# Tonically active interneurons gate motor output in *Drosophila* larvae

**DOI:** 10.64898/2026.01.19.700243

**Authors:** Takahisa Date, Yingtao Liu, Akihiro Yamaguchi, Paul McNulty, Rui Wu, Marc Gershow, Akinao Nose, Maarten F Zwart, Hiroshi Kohsaka

## Abstract

Tonic inhibition and its transient removal are fundamental neural mechanisms for gating activity, suppressing out-of-phase excitation, and enhancing the temporal precision of behavior. However, their roles within locomotor central pattern generators (CPGs) remain poorly understood. Here, we identify A02l neurons—segmentally repeated excitatory interneurons in the *Drosophila* larval ventral nerve cord (VNC)—that regulate crawling behavior through tonic inhibition. A02l neurons exhibit high baseline activity and are transiently suppressed during motor output. They provide excitatory input to inhibitory premotor networks, enabling widespread silencing of motor neurons to maintain muscle relaxation. Optogenetic activation of A02l neurons reduced crawling frequency, whereas their inhibition induced transient whole-body contractions. Together, these findings reveal a cellular-resolution mechanism for tonic inhibition within the nerve cord and highlight its importance in gating motor output and maintaining motor quiescence.

## Introduction

Inhibitory synaptic input plays a central role in shaping the spatiotemporal patterns of motor output. In many motor systems, inhibition is predominantly phasic and recruited by phasic excitation. Classic examples include recurrent inhibition of motoneurons via Renshaw cells, reciprocal suppression of antagonistic muscles through Ia afferent feedback during contraction ^1,2^, and left-right alternation driven by cross-inhibitory interactions triggered by unilateral excitation ^3,4^. These mechanisms position phasic inhibition as a primary mode of shaping motor patterns.

In contrast, tonic inhibition regulates neuronal excitability and behavioral selection. In cat ankle extensor systems, tonic Ia afferent input modulates motoneuron excitability and force output ^5^; in crayfish, tonic inhibition of the lateral giant fibers reduces the probability of escape behavior ^6^; and in the vertebrate basal ganglia, tonic firing mediates behavioral selection ^7,8^. Despite these examples highlighting the importance of tonic inhibition in selecting actions, how tonic inhibition contributes to shaping patterned motor output within the locomotor CPG itself remains poorly understood.

Addressing this gap requires examining rhythmic motor circuits at the cellular and circuit levels. Axial locomotion is a widespread mode of animal movement ^9^. It is generated by the propagation of neural activity along the anterior-posterior axis, which is translated into coordinated muscle contraction during behaviors such as crawling, swimming, and walking. Core components of segmental motor circuits have been identified across diverse taxa ^10^ – including leech ^11^, nematode ^12^, fly larva ^13,14^, zebrafish ^15^, tadpole ^16^, and mouse ^4^. Within these circuits, phasic inhibition plays critical roles in both intra-segmental and inter-segmental coordination of locomotion CPGs. By contrast, although evidence exists for tonically active premotor interneurons ^17–21^, the specific cell types involved and the functional roles of tonic inhibition within locomotor CPGs remain poorly understood.

*Drosophila* larvae have emerged as a powerful model for investigating neural circuits underlying axial locomotion. Their bodies comprise repeated segments (**Figure 1A**), and locomotion is generated by the central nervous system, consisting of the brain and ventral nerve cord (VNC), which functionally parallels the vertebrate spinal cord. Contraction of each body-wall segment is controlled by neuromeres, which are repeated segmental neural circuit units within the VNC (**Figure 1B**). Larvae move forward or backward through sequential segmental muscle contractions, and the VNC can produce propagating activity waves in *ex vivo* preparations even in the absence of sensory feedback ^22^, demonstrating an intrinsic CPG.

**Figure 1.**
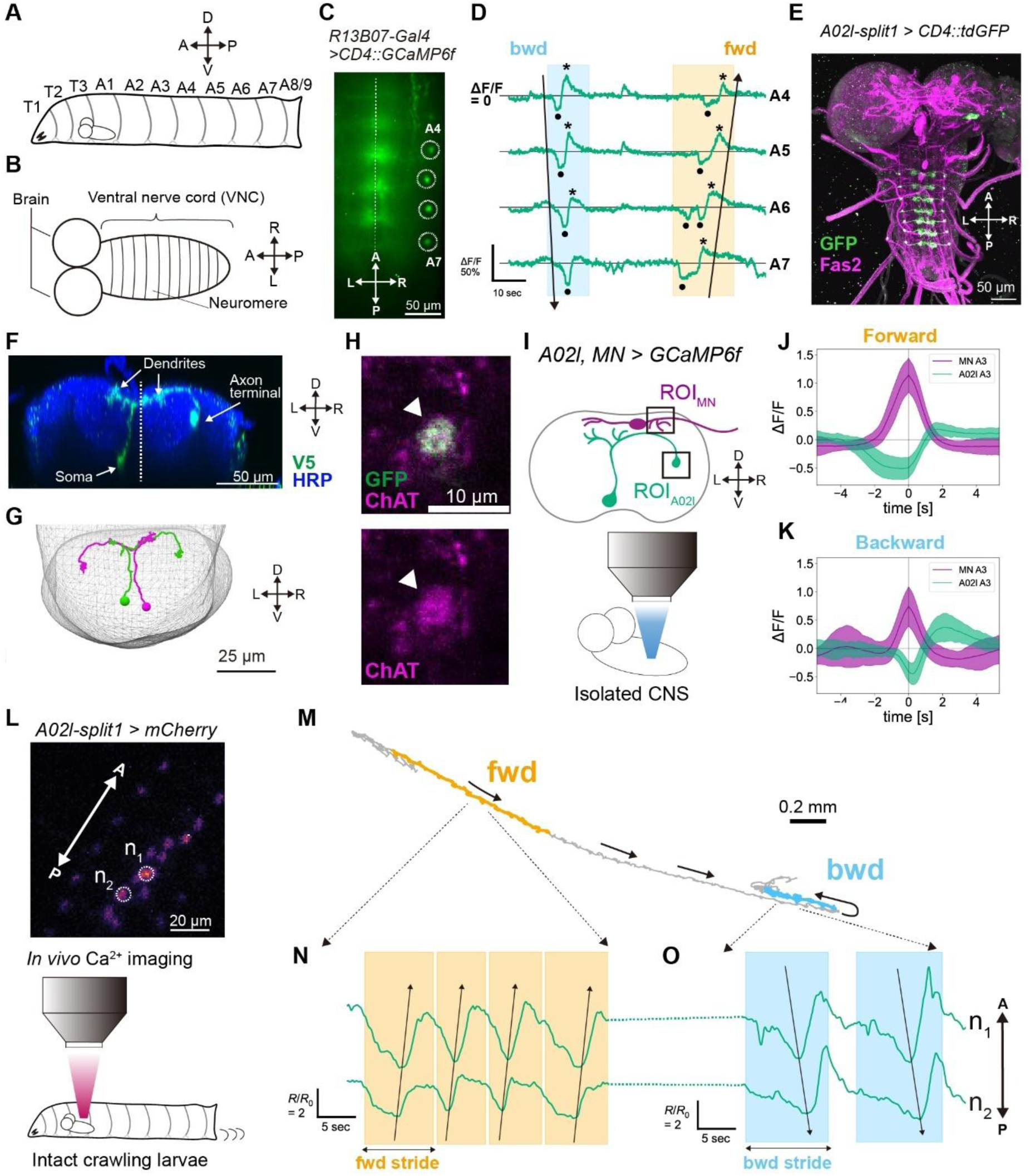
Segmental A02l interneurons in the Drosophila larval ventral nerve cord show calcium dips during motor output. **(A)** Schematic of *Drosophila* larval body organization. Larvae consist of three thoracic segments (T1-T3) and eight abdominal segments (A1–A8/9). The central nervous system (CNS) is in the T3-A1 segments. A, anterior; P, posterior; D, dorsal; V, ventral. **(B)** Schematic of the larval central nervous system (CNS), comprising the brain and the ventral nerve cord (VNC). The VNC consists of segmentally repeated neuromeres corresponding to body segments. A, anterior; P, posterior; L, left; R, right. **(C)** Representative calcium imaging frame from *R13B07-Gal4 > CD4::GCaMP6f* larvae. Dashed circles indicate spherical structures in the A4–A7 neuromeres; the dashed line marks the VNC midline. A, anterior; P, posterior; L, left; R, right. **(D)** Representative calcium traces from the regions of interest shown in (C). One backward (bwd) and one forward (fwd) fictive waves are indicated by arrows and shaded bands. Horizontal lines indicate *Δf*/*f* = *0* for each neuron. Black dots denote calcium dips; asterisks indicate rebound events. **(E)** Expression pattern of A02l-split1 (*R84G07-AD; R95H02-DBD > CD4::tdGFP*), labeling A02l neurons in A1-A7 neuromeres. A, anterior; P, posterior; L, left; R, right. **(F)** Top: Morphology of a single A02l neuron visualized using the Multi-Color FlipOut method ^38^ (*R95H02-Gal4>MCFO4*). The dashed line is the midline of the VNC. The soma, axon terminal, and dendrites are indicated by arrows. D, dorsal; V, ventral; L, left; R, right. **(G)** A02l neurons in the A1 neuromere reconstructed from EM connectomics data. Green, left A02l neuron; magenta, right A02l neuron. D, dorsal; V, ventral; L, left; R, right. **(H)** A02l axon terminals co-stained with choline acetyltransferase (ChAT). Green, A02l axon terminals (*A02l-split1 > CD4::GCaMP6f*); magenta, ChAT. **(I)** Experimental setup for *ex vivo* calcium imaging of A02l neurons and motor neurons. GCaMP6f was expressed in A02l neurons and aCC motor neurons (*A02l-split, eve-Gal4 > CD4::GCaMP6f*). Signals were extracted from motor neuron dendrites (ROIMN) and A02l axon terminals (ROIA02l). **(J, K)** Averaged ΔF/F traces of A02l neurons and motor neurons in the A3 segment during fictive forward waves (J; 94 calcium-dip events from 8 animals) and backward waves (K; 52 events from 5 animals). Time 0 indicates the motor neuron activity peak. Shaded regions indicate standard deviation. **(L)** Schematic of in vivo calcium imaging in intact crawling larvae. Two-photon imaging was performed while tracking A02l neurons. In the example shown in (**M–O**), two A02l somata (n1 and n2) were tracked. Genotype: *A02l-split1 > 6XmCherry, GCaMP6f*. A, anterior; P, posterior. **(M)** Larval trajectory during in vivo calcium imaging. The larva transitioned from forward to backward crawling. **(N, O)** Calcium traces from two A02l somata during forward (**N**) or backward (**O**) crawling. Signals are expressed as *R* = *f*_*GCaMP*_/*f_mCherry_*, normalized to the minimum value (*R_0_*, see methods). Both neurons were transiently inhibited during each crawling event (shaded bar). Calcium dips propagate in the direction of locomotion. A, anterior; P, posterior.

Cell-type-specific driver lines ^23–25^ and EM connectomics ^26,27^ have enabled the identification of premotor neural circuits essential for larval locomotion ^13,28,29^. Current models posit that motor neurons receive transient excitation to trigger contraction, together with phase-specific inhibition that can precede or follow excitation depending on wave direction and segmental context. For example, during forward waves, the cholinergic neuron A27h provides excitatory synaptic input to the GABAergic neuron GDL in the next anterior segment to prevent premature contraction ^30^. The cholinergic neurons Ifb-Fwd and Ifb-Bwd are selectively active during forward and backward waves, respectively, and deliver excitatory synaptic input to inhibitory premotor interneurons in adjacent segments that have already contracted ^31^. All VNC interneurons described to date exhibit transient, phase-specific activity. To our knowledge, sustained inhibition has not been implicated in segmental control during larval axial locomotion.

Here, we identify A02l neurons as a segmentally-repeated class of cholinergic interneurons in the larval VNC that exhibit tonic activity during rest and transient suppression with rebound during motor output. These dynamics are anti-correlated with motor neuron activity. Simultaneous calcium and voltage imaging shows that the suppression coincides with hyperpolarization, indicating that A02l neurons are continuously active and transiently inhibited during wave propagation. Connectomic analysis reveals that A02l neurons target inhibitory premotor interneurons that suppress longitudinal muscle contraction, suggesting that A02l contributes to maintaining a segmental resting state. Consistent with this, optogenetic activation of A02l neurons suppresses larval locomotion, whereas their inhibition induces muscle contraction. Together, these findings suggest that the larval VNC houses a circuit dedicated to suppressing involuntary muscle activation through tonic inhibition.

## Results

### A02l neurons exhibit tonic activity with transient suppression during larval motor output

To characterize the activity patterns of interneurons in *Drosophila* larvae, we expressed the calcium indicator GCaMP6f ^32^ using Gal4 lines that label sparse VNC interneuron populations and performed calcium imaging in an *ex vivo* preparation ^33^. Among these, R13B07-Gal4 labeled segmentally repeated spherical structures located on the lateral side of the neuropil (**Figure 1C**) that displayed (1) high baseline GCaMP fluorescence, suggesting tonic activation, (2) transient decreases (“dips”), and (3) subsequent rebound activity (**Figure 1D**). To our knowledge, no previously described interneuron type in larval motor circuits exhibits this set of activity patterns. R13B07-Gal4 was previously reported to label A02l neurons, segmental interneurons with clustered axon terminals ^27^. These neurons were characterized in EM connectomics analysis as postsynaptic targets of dbd sensory neurons ^27^ and presynaptic partners of A08a interneurons ^34,35^, but their role in larval locomotion has not been examined. Based on their characteristic spherical morphology observed in calcium imaging and their anatomical location, we conclude that the neuron exhibiting calcium dips in the R13B07-Gal4 line corresponds to A02l.

To refine cell-type specificity, we visually screened Gal4 expression patterns from the Janelia FlyLight database ^24^ for the characteristic A02l axon terminal-like structures. We then identified and generated several Gal4 and split-Gal4 lines that selectively label A02l neurons (**Figures 1E, S1A, and S1B**). The cell bodies of A02l reside ventrally in the VNC, with primary neurites projecting dorsally and forming tightly clustered presynaptic terminals on the contralateral side of the neuropil (**Figure 1F**), consistent with the reconstructed A02l morphology in the connectomics data (**Figure 1G**). Immunostaining confirmed that the terminals are cholinergic, indicating that A02l provides excitatory outputs (**Figures 1H, S1C, and S1D**).

Using the A02l-split1 Gal4 line, which selectively labels A02l neurons in neuromeres A1-A7 (**Figure 1E**), we examined A02l activity in *ex vivo* preparations without contributions from other R13B07-Gal4-positive abdominal neurons. A02l axon terminals in all abdominal neuromeres show calcium dips and rebounds propagating forward or backward (**Figures S1E and S1F, Movie S1**). This observation implies that the activity of A02l is coupled with segmental motor activity during peristaltic locomotion. To test this possibility, we performed calcium imaging of A02l neurons and motor neurons simultaneously in the *ex vivo* preparation (**Figure 1I**). Motor neurons exhibit transient, propagating activation that reflects fictive peristaltic locomotion ^36^. Consistent with our hypothesis, A02l dips and rebounds coincided with motoneuronal activation (**Figures 1J, 1K, S2A, and S2B; Movie S2**). In addition, A02l neurons showed significantly larger rebound activity during the backward waves (*Δf*/*f* rebound peak amplitude in forward wave: 0.273 ± 0.110, backward wave: 0.471 ± 0.238, p = 1.1 × 10^-9^ with Mann-Whitney U-test).

We further examined A02l activity in intact moving larvae using a neuron-tracking microscope equipped with acousto-optic deflectors ^37^ (**Figure 1L**). Consistent with calcium imaging in *ex vivo* preparations, A02l somata showed transient suppression in both forward and backward locomotor directions (**Figures 1M-1O**, **Figures S2C-S2F, Movie S3**).

Taken together, these results suggest that A02l neurons are tonically active and exhibit transient suppression and rebound coupled to segmental motor output during peristaltic locomotion.

### Membrane potential contributes to calcium dynamics in A02l neurons

We next examined how membrane potential shapes the calcium dynamics of A02l neurons. To this end, we performed simultaneous dual-color calcium and voltage imaging of A02l axon terminals in *ex vivo* preparations, using the red calcium indicator jRGECO1a ^39^ and the green voltage sensor ASAP5 ^40^ (**Figures 2A**). When calcium signals at A02l presynaptic terminals transiently decreased, membrane voltage simultaneously exhibited hyperpolarization (**Figures 2B and 2C**). Notably, voltage signals did not show positive deflections corresponding to the rebound phase of the calcium signal. These observations indicate that membrane-potential changes do not account for the rebound dynamics but do underlie dip formation.

**Figure 2.**
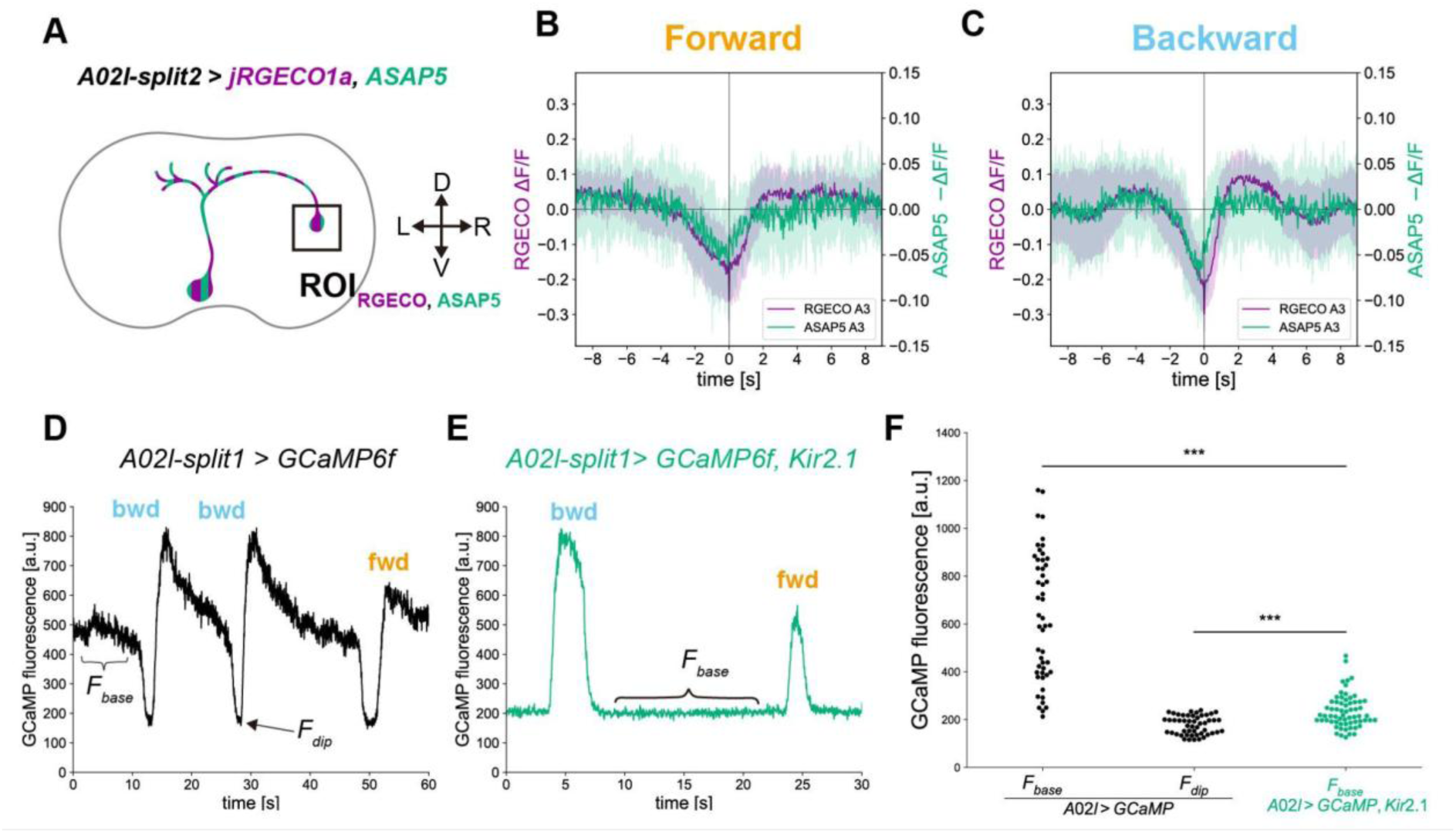
A02l neurons hyperpolarize during calcium dips. **(A)** Schematic indicating the regions of interest (ROIs) for simultaneous calcium and voltage imaging in A02l neurons. jRGECO1a and ASAP5 were expressed in A02l neurons, and signals were analyzed in axon terminals. **(B, C)** Averaged ΔF/F traces of jRGECO1a (magenta) and ASAP5 signals (green) from A02l neurons in the A3 neuromere during forward waves (**B**; 19 calcium-dip events from 5 animals) and backward waves (**C**, 28 events from 4 animals). For ASAP5, the sign-inverted ΔF/F is plotted to represent membrane potential changes. Time 0 corresponds to the negative peak of A02l activity. Shaded bands indicate standard deviation. **(D)** Representative *ex vivo* calcium traces from an A02l axon terminal in the A3 neuromere (*A02l-split1 > CD4::GCaMP6f*), showing calcium dips. The time points used to define baseline fluorescence (*f_base_*) and dip fluorescence (*f_dip_*) are indicated. **(E)** Representative *ex vivo* calcium traces from an A02l axon terminal in the A3 neuromere expressing Kir2.1 (*A02l-split1 > CD4::GCaMP6f, Kir2.1::GFP*), which show only positive transients. The time window used to define baseline fluorescence (*f_base_*) is indicated. **(F)** Quantification of baseline GCaMP fluorescence (*f_base_*) and dip fluorescence (*f_dip_*) using the criteria illustrated by representative examples (D) and (E). Control neurons (*A02l > GCaMP*; n=50 cells from 5 animals) are shown in black, and Kir2.1-expressing neurons (*A02l > GCaMP, Kir2.1*; n=64 cells from 5 animals) are shown in green. Kir2.1 expression was significantly reduced *f_base_* to levels comparable to *f_dip_* in control neurons. Brunner-Munzel test with Holm-Bonferroni correction. *** p < 0.001.

To further test the role of membrane potential, we analyzed calcium signals in A02l neurons while hyperpolarizing them with Kir2.1 ^41^, a modified inward-rectifier potassium channel. Hyperpolarized Kir2.1-expressing A02l neurons no longer exhibited calcium dips, but instead only showed transient increases in calcium signals (**Figures 2D and 2E**). The baseline fluorescence of Kir2.1-expressing A02l neurons is significantly lower than that of the control group, comparable to the fluorescence in calcium dips under control conditions (**Figure 2F**). This finding is consistent with the voltage imaging results: calcium dips coincide with hyperpolarization, whereas calcium rebounds are not accompanied by depolarization. Notably, the transient calcium increases occurred independently of the dips, indicating that they are unlikely to arise from rebound depolarization. Taken together, these results suggest that the calcium dips in A02l neurons arise from hyperpolarization of the membrane potential, likely driven by synaptic inputs from upstream inhibitory partners.

### Coordinated neurotransmitter dynamics accompany transient suppression of A02l neurons

We hypothesized that the transient hyperpolarization of A02l neurons is driven by the inhibitory inputs on A02l neurons. To verify the existence of the inhibitory input, we performed simultaneous calcium and GABA imaging of A02l neurons using the red calcium indicator jRGECO1b ^39^ and the green GABA sensor iGABASnFR2 ^42^ (**Figure 3A**). GABA signals in the vicinity of A02l dendrites increased at the timing of calcium dips during fictive crawling (**Figures 3B and 3C**), indicating that dendritic GABAergic inhibition contributed to the hyperpolarization of A02l neurons. Low-amplitude GABA signals were also detected at axon terminals during motor output, suggesting axo-axonic inhibitory modulation (Supplementary **Figures S3A-S3C**).

**Figure 3.**
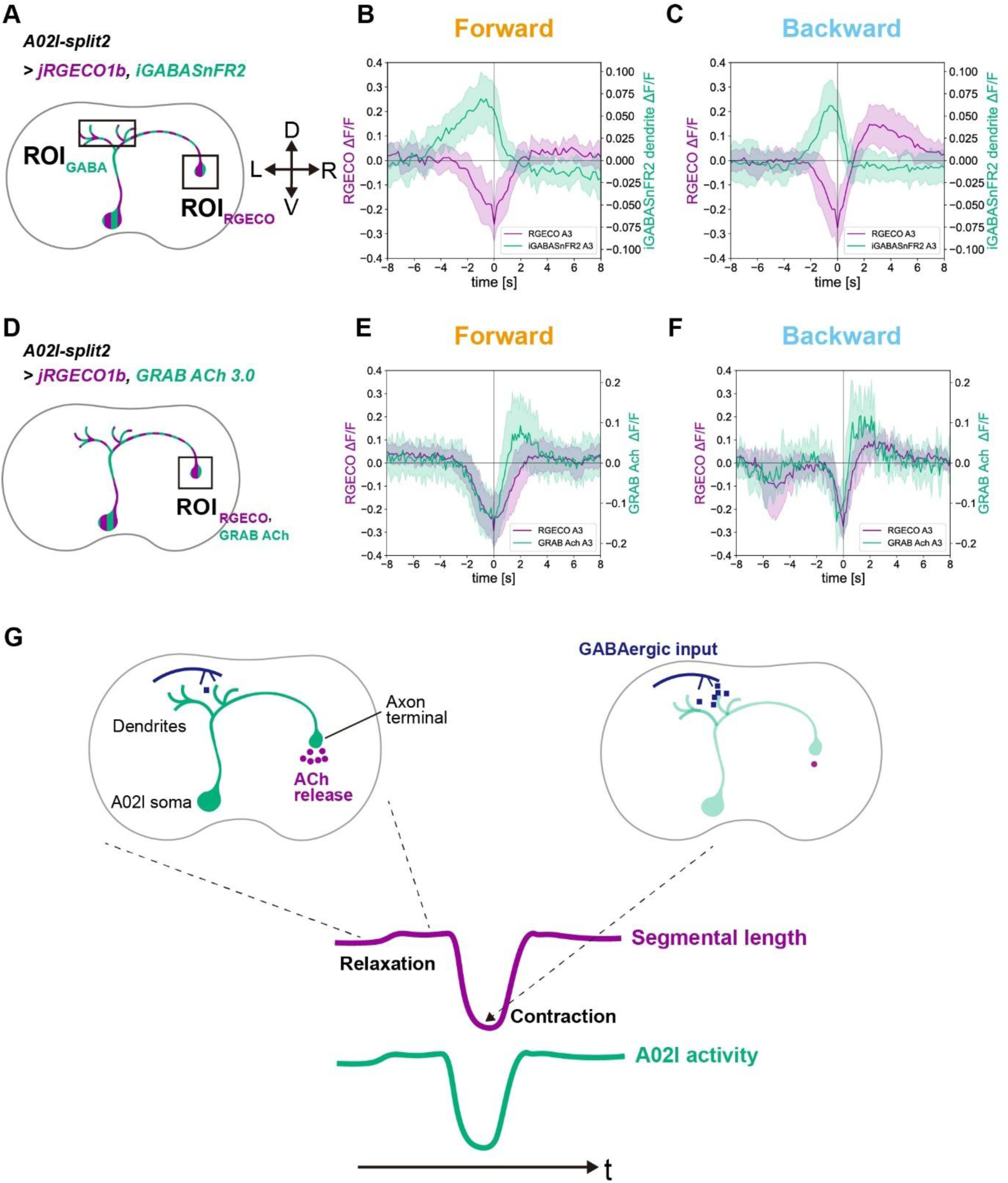
Time course of GABA reception and acetylcholine release in A02l neurons. **(A)** Schematic indicating the ROIs for simultaneous calcium and GABA imaging in A02l neurons. jRGECO1b was analyzed in axon terminals, and iGABASnFR2 was analyzed in dendrites (*A02l-split2>jRGECO1b, iGABASnFR2*). **(B, C)** Averaged ΔF/F traces of jRGECO1b in axon terminals (magenta) and iGABASnFR2 in dendrites (green) from A02l neurons in the A3 neuromere during forward waves (**B**, 17 calcium-dip events from 4 animals) and backward waves (**C**, 18 events from 6 animals) Time 0 corresponds to the negative peak in A02l activity. Shaded bands indicate standard deviation. **(D)** Schematic indicating the ROIs for simultaneous calcium and acetylcholine imaging in A02l neurons. jRGECO1b and GRAB-ACh3.0 were expressed in A02l neurons, and both signals were analyzed in axon terminals (*A02l-split2>jRGECO1b, GRAB-ACh3.0*). **(E, F)** Averaged ΔF/F traces of jRGECO1b (magenta) and GRAB-ACh3.0 (green) from A02l axon terminals in the A3 neuromere during forward waves (**E**, 25 events from 8 animals) and backward waves (**F**, 9 events from 4 animals). Time 0 corresponds to the negative peak of A02l activity. Shaded bands indicate standard deviation. **(G)** Schematic illustrating A02l dynamics and segmental length. The bottom traces show segmental length (magenta) and A02l activity (green). When the segment is relaxed, the A02l neuron maintains a high membrane potential and tonically releases acetylcholine (upper left). During segmental contraction, GABAergic input to the A02l dendrites hyperpolarizes the neuron during motor output in the corresponding segment, thereby suppressing acetylcholine release (upper right). D, dorsal; V, ventral; L, left; R, right.

To evaluate excitatory signaling, we conducted simultaneous calcium and acetylcholine imaging of A02l neurons using the red calcium indicator jRGECO1b and the green acetylcholine sensor GRAB-ACh3.0 ^43^ (**Figure 3D**). Acetylcholine signals near A02l axon terminals exhibited transient dips and rebounds that mirrored A02l calcium dynamics (**Figure 3E and 3F**). The transient suppression of acetylcholine suggests that A02l neurons normally exhibit baseline acetylcholine release during rest, and reduce release during motor output.

We next investigated how A02l neurons maintain their tonic activity in the resting state. To assess the contribution of A02l neuron spiking and their action potential-mediated presynaptic drive, we blocked voltage-gated sodium channels with tetrodotoxin (TTX) in *ex vivo* preparations and recorded calcium signals before and after drug applications (**Figure S4**). After applying TTX, basal GCaMP signals decreased (**Figure S4C**), indicating that presynaptic spiking and/or spiking of A02l contribute to maintaining the tonic activity of A02l. In saline, A02l neurons exhibit calcium dips, whose amplitudes could reflect GABA-mediated hyperpolarisation (**Figure S4B**). When we compared these dips in saline with the basal GCaMP signal level under TTX, the latter is consistently higher **(Figure S4D**). This indicates that A02l neurons maintain residual tonic activity even in the absence of spiking, or under conditions in which spiking is strongly suppressed. This reveals a spike-independent mechanism underlying tonic A02l activity in the resting state, and could even indicate A02l is non-spiking. Together, these results indicate that A02l neurons are tonically active during rest and continuously release acetylcholine (**Figure 3G**). During motor output, dendritic GABAergic inhibition transiently hyperpolarizes A02l neurons, leading to temporary reductions in intracellular calcium and acetylcholine release.

### A02l neurons integrate premotor activity and proprioceptive feedback and target inhibitory first-order premotor interneurons

To identify the circuit mechanisms underlying A02l activity, we examined their synaptic partners using an electron microscopy (EM)-based connectomics dataset ^26,27^. We investigated the upstream and downstream connections of two A02l neurons in the A1 neuromere and identified presynaptic and postsynaptic partners (**Figures 4A-4D**, **Tables S1 and S2**).

**Figure 4:**
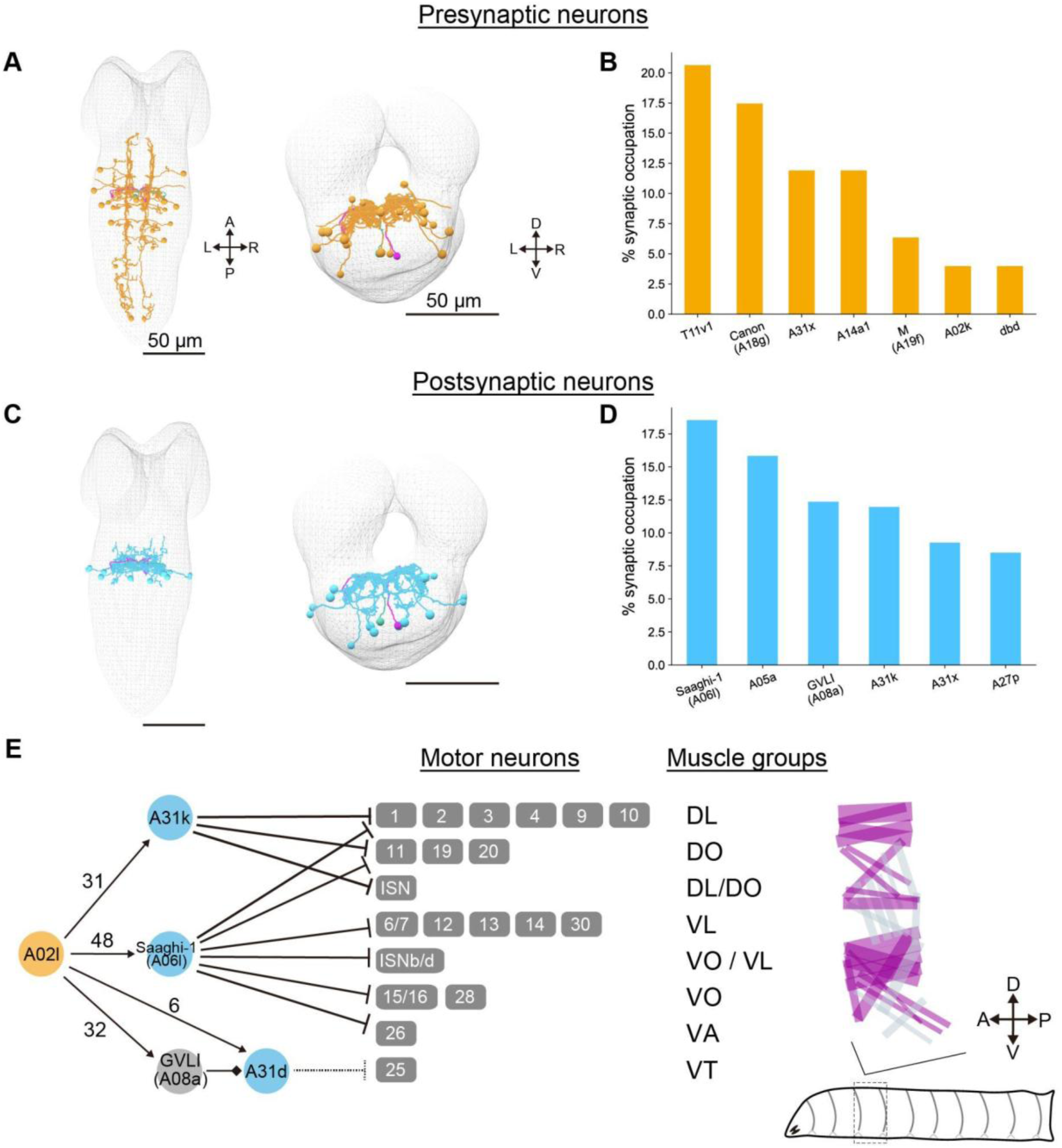
Connectomics analysis revealed that A02l neurons are a part of the muscle-relaxing circuit. **(A)** Reconstruction of major presynaptic partners of A02l neurons in the A1 neuromere. Left, dorsal view; right, frontal view. A, anterior; P, posterior; D, dorsal; V, ventral; L, left; R, right. **(B)** Percentage of total synaptic inputs to A02l neurons in the A1 neuromere. Major presynaptic partners (≥ 3.5% of total synapses) are shown. **(C)** Reconstructions of major postsynaptic partners of A02l neurons in the A1 neuromere. Left, dorsal view; right, frontal view. **(D)** Percentage of total synaptic outputs from A02l neuron in the A1 neuromere. Major postsynaptic partners (≥ 3.5% of total synapses) are shown. **(E)** Downstream circuit diagram of A02l neurons. Nodes represent left-right pairs of homologous neurons within a neuromere. The cholinergic excitatory A02l neuron is shown in orange; GABAergic inhibitory neurons are shown in cyan. Arrows indicate excitatory connections, bar-headed lines indicate inhibitory connections, and diamond-headed lines indicate glutamatergic synapses. Numbers adjacent to edges indicate the number of synaptic contacts. Putative GABAergic A31d connections are indicated by dashed lines. Motor neurons (MNs) and their target muscles are shown centrally. The muscle map is displayed on the right side, with muscles inhibited by A02l downstream neurons shown in magenta. Scale bars, 50 μm.

First, we identified seven major presynaptic partners: (1) local interneurons A14a1, A31x, and A02k, (2) abdominal ascending neurons Canon (A18g), (3) abdominal ascending neurons M (A19f), (4) thoracic descending neurons T11v1, and (5) dorsal bipolar dendritic (dbd) sensory neurons (**Figures 4A and 4B**).

A14a1 neurons accounted for 11% of total presynaptic inputs (**Figure S5E**). A14a1 neurons are derived from neuroblast lineage 14, which gives rise to GABAergic interneurons, including A14a neurons ^44^. These have a similar morphology to A14a1, suggesting that A14a1 provides inhibitory input to A02l neurons. This inference is consistent with our GABA imaging, which showed increased GABA signals at calcium dips (**Figures 3A–3C**), implicating A14a1 as a candidate source of dip-triggering inhibition. A31x neurons are segmental interneurons that form reciprocal axo-axonic connections with A02l neurons (**Figure S5D**), whereas A02k neurons, which arise from the same neuroblast as A02l neurons, form axo-dendritic synapses onto A02l (**Figure S5G**).

There are three classes of descending/ascending neurons innervating A02l neurons. Canon neurons (A18g, **Figure S5B**) are excitatory neurons activated during backward, but not forward, locomotion ^45^. The greater A02l rebound amplitude during backward than forward waves (**Figures 1J and 1K**) is therefore likely mediated by the backward wave-specific activity of Canon neurons (**Figure S5B**). M (A19f) neurons are ascending interneurons in the abdominal segments (**Figure S5F**) implicated in the emergence of embryonic locomotor waves ^46^ and in larval locomotion ^47^. T11v1 neurons are descending interneurons located in neuromeres T1–T3 and target A02l neurons in abdominal neuromeres (**Figure S5A**). These descending/ascending inputs may control intersegmental coordination of A02l activity.

Dbd sensory neurons are proprioceptive stretch receptors in the larval body wall ^48^ (**Figure S5H**). Dbd activity decreases during longitudinal muscle contraction, displaying an anti-correlated pattern with motor output ^49^, similar to A02l calcium dynamics (**Figure S5I**).

Taken together, these presynaptic neuronal classes indicate that A02l neurons integrate proprioceptive afferents, intrasegmental inhibition, and intersegmental excitatory inputs to generate their characteristic, phase-timed calcium dynamics.

To determine how A02l neurons influence motor output, we next analyzed their downstream connectivity in the A1 neuromere. Most postsynaptic partners were located within the same A1 neuromere (38/45 neurons), indicating that each A02l neuron regulates motor output segmentally. The postsynaptic partners consist of six main neuron types (**Figures 4C and 4D).** Notably, A02l neurons make synapses onto Saaghi-1 (A06l), GVLI(A08a), and A31k neurons (**Figure S6**), all of which are direct or indirect inhibitory premotor neurons ^50–52^. Previous studies have shown that optogenetic activation of GVLI or A31k neurons halts larval locomotion ^45,51^. These downstream neurons collectively regulate 20 of the 30 larval body-wall muscles (**Figure 4E**), including all longitudinal muscles, the principal drivers of axial locomotion ^33^. These data indicate that A02l neurons are positioned to promote muscle relaxation by engaging inhibitory premotor circuits through different segmental outputs that influence multiple muscles within each segment.

The connectomics analysis above indicates that A02l integrates synaptic inputs from diverse upstream interneurons and sensory neurons (Pre-A02l) and distributes divergent outputs to inhibitory premotor circuits (Post-A02l). The circuit diagram, featuring a large degree of interconnectivity, shows that A02l does not necessarily act as a bottleneck (a single mandatory relay), but rather as a core node; it is one of several parallel elements in the information flow from Pre-A02l to Post-A02l neurons (**Figure S6F**). These observations suggest that A02l provides a global tonic drive to inhibitory premotor neurons, shaping motor output under the control of multiple upstream pathways.

### Optogenetic activation of A02l neurons suppresses larval locomotion, whereas inhibition of A02l neurons evokes muscle contraction

Because A02l neurons are tonically active during periods without motor output and connect to premotor inhibitory circuits, we hypothesized that A02l activity maintains a resting motor state of the segment. To test this, we optogenetically activated A02l neurons by expressing CsChrimson ^53^ specifically in A02l neurons (**Figure 5A**) and activated them using 660 nm light. Photoactivation halted locomotion and significantly reduced crawling frequency (**Figures 5B-5G, Movie S4**), indicating that A02l activity suppresses motor output.

**Figure 5.**
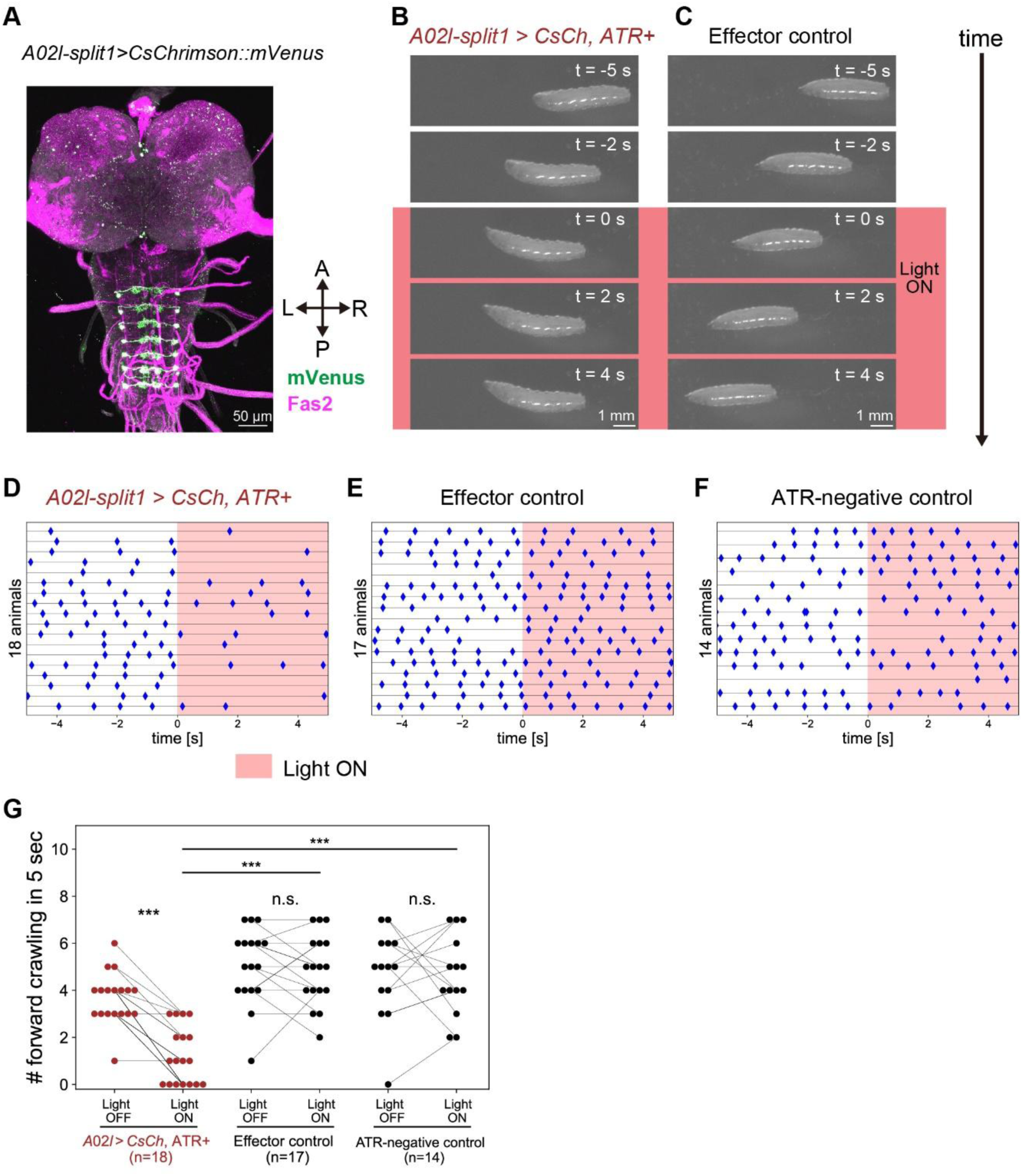
Optogenetic activation of A02l neurons inhibits larval crawling. **(A)** Expression pattern of CsChrimson::mVenus (CsCh) driven by *A02l-split1 Gal4*. A, anterior; P, posterior; L, left; R, right. **(B, C)** Representative behavior of larvae in the experimental group (*A02l-split1 > CsCh, ATR+*) and the effector control group (*+/UAS-CsCh, ATR+*), respectively. Time 0 indicates the onset of red-light stimulation (red background). The larva in the experimental group **(B)** ceases crawling during photostimulation (t = 0 - 4 s). **(D-F)** Raster plots of forward crawling events in the experimental group (**D**), effector control group (**E**), and ATR-negative control group (**F**). Crawling initiation (onset of tail contraction) is indicated by blue diamond symbols. Each row shows a 10-second recording from an individual larva (n = 18 in **D**, n = 17 in **E**, and n = 14 in panel **F**). The five-second light stimulation period is shown with a red bar. **(G)** Quantification of the number of forward crawling events before and during photostimulation. Effector control: *+/UAS-CsCh*, ATR-fed; ATR-negative control: *A02l-split1 > CsCh*, non-ATR-fed. Within-group comparisons between light OFF and ON periods were performed using the Wilcoxon signed-rank test. Comparisons across genotypes during stimulation were performed using the Mann-Whitney U test with Holm-Bonferroni correction. *** p < 0.001.

As A02l-split1 occasionally labelled dbd sensory neurons (**Figure S7A**), we assessed possible off-target contribution to the phenotype. Optogenetic activation of single dbd neurons (**Figure S7B**) using a heat-shock flip-out technique did not affect crawling frequency (**Figure S7C**). In addition, photoactivation using A02l-split2 Gal4, which labels a subset of A02l neurons without off-target expression, similarly reduced locomotor activity (**Figures S7D and S7E**). These controls confirm that activation of A02l neurons, rather than dbd neurons, suppresses locomotion.

In a complementary set of experiments, to examine the role of A02l activity in maintaining segmental relaxation, we optogenetically inhibited A02l neurons using GtACR1 ^54,55^ driven by A02l-split2 Gal4 (**Figure 6A**). Photoinhibition of A02l neurons induced transient whole-body contractions (**Figures 6B and 6C, Movie S5**), displayed as a significant decrease in body-wall area relative to effector and ATR-negative controls (**Figures 6D and 6E**). Together, these bidirectional manipulations demonstrate that A02l activity suppresses muscle contraction and is required to maintain the segmental resting motor state.

**Figure 6.**
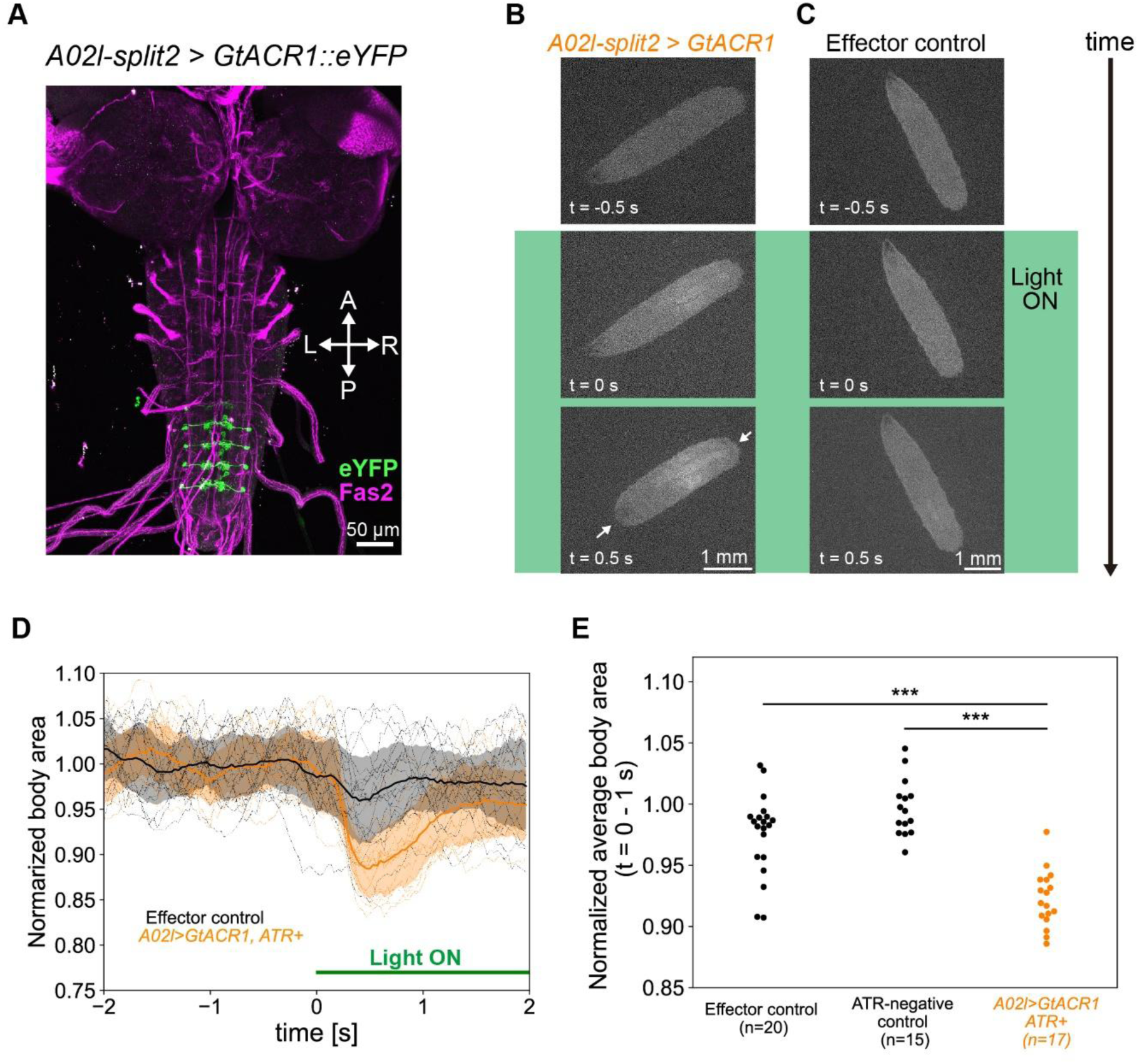
Optogenetic inhibition of A02l neurons induces a transient whole-body contraction. **(A)** Expression pattern of GtACR1 driven by *A02l-split2*. A02l neurons in A4 –A7 neuromeres express GtACR1. Green: eYFP; magenta: Fas2. A, anterior; P, posterior; L, left; R, right. **(B, C)** Representative behavior of larvae in the experimental group (**B**; *A02l-split2 > GtACR1, ATR+*) and the effector control group (**C**; *+/ UAS-GtACR1, ATR+*). Time 0 indicates the onset of green-light stimulation. The larval body rapidly contracts following photostimulation in the experimental group but not in the control group. **(D)** Time course of normalized body area in the experimental group (*A02l-split2 > GtACR1*, ATR+, orange traces) and the effector control group (*+/UAS-GtACR1*, ATR+, black traces). Individual larvae are shown as dotted lines; mean traces as solid lines; and standard deviation as shaded bands. Green light was applied from t = 0 to 2 s. Body area was normalized to the average value during the pre-stimulation period (t = -2 to 0 s). **(E)** Comparison of the normalized body area during photostimulation (t = 0 to 1 s) among the effector control group (n = 20 animals), ATR-negative control group (n = 15 animals), and experimental group (in orange, n=17 animals). Mann-Whitney U test followed by Holm-Bonferroni correction. ***p < 0.001.

## Discussion

In animal motor control, maintaining muscle relaxation is as critical as driving muscle contraction. Previous studies have established that phasic inhibition within the CPGs plays a key role in shaping coordinated motor output by timing muscle activation and enforcing appropriate phase relationships ^1,56–58^. However, whether tonic inhibition contributes to the maintenance of muscle relaxation and the gating of motor output within premotor CPG networks remains unclear. In this paper, we identify A02l neurons as a key component of this mechanism. A02l neurons exhibit calcium activity that is anti-correlated with motor neuron activity, and optogenetic activation or inhibition of A02l neurons halts locomotion or induces whole-body contraction, respectively. These results indicate that A02l neurons form a critical circuit module that maintains muscle relaxation and gates motor output. By suppressing involuntary activity while selectively permitting motor commands, this module is likely to enhance the robustness of locomotor control (**Figure 7**).

**Figure 7.**
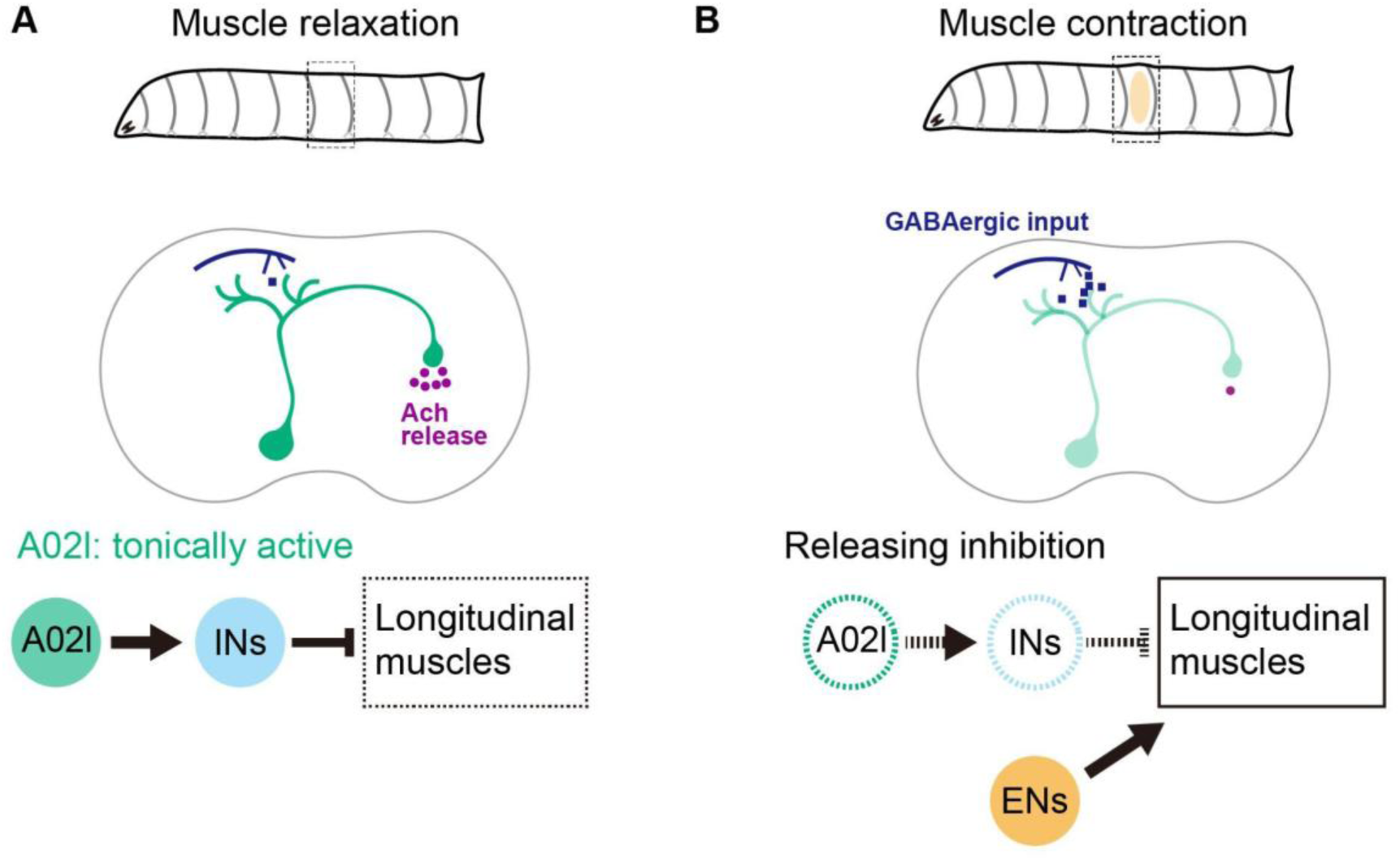
Gating of motor output with A02l neurons. **(A)** When a segment is relaxed (top), A02l neurons in the corresponding neuromere are tonically active (middle) and maintain muscle relaxation. A02l neurons activate inhibitory interneurons (INs), thereby suppressing longitudinal muscle activity (bottom). **(B)** During longitudinal muscle contraction during crawling (top), A02l neurons are transiently inhibited (middle). This transient inhibition gates motor output by releasing tonic inhibition, allowing other excitatory interneurons (ENs) to drive muscle contraction.

### Tonic activity in A02l neurons

Our study identifies A02l neurons as a class of interneurons that exhibits tonic calcium activity maintained by membrane depolarization, as revealed by Kir2.1-mediated hyperpolarization experiments. Similar forms of tonic activity have been reported in other motor systems. For example, in the cat spinal cord, motoneurons receive coordinated tonic excitation and inhibition ^5^, and in *C. elegans*, AVA interneurons maintain a depolarized membrane potential sufficient to drive continuous vesicle release ^17^. Consistent with these observations, our imaging data suggest continuous acetylcholine release from A02l neurons during periods without segmental contraction.

Sustained neurotransmitter release is metabolically costly ^59–63^, requiring ongoing vesicle loading, transport, fusion, and recycling, and most importantly, Na^+^/K^+^-ATPase activity to maintain ion gradients. The relative rarity of tonic neurons in the nervous system may therefore reflect these energetic constraints, implying that tonic signaling is employed selectively where it serves critical circuit functions. In spiking neurons, tonic activity can be generated by multiple molecular mechanisms, including persistent sodium currents ^64^, low-threshold L-type calcium channels ^65^, sodium leak channels ^66^, and second-messenger-dependent modulation of ion channels ^67^. Persistent activity is also observed in non-spiking neurons in invertebrates, where tonic depolarization supports graded neurotransmitter release in motor circuits, such as in the locust ^68,69^ and nematode ^17,70^.

The molecular and circuit mechanisms that maintain tonic depolarization in A02l remain unknown, as is whether A02l is a spiking neuron. Answering these questions may provide broader insight into how neural circuits sustain baseline activity states and actively regulate motor output.

### Roles of A02l neurons in motor control

Connectomics and optogenetic analyses demonstrate that A02l neurons continuously suppress muscle contraction via inhibitory premotor circuits, while transient dips in A02l activity permit segmental contraction. One potential function of this architecture is noise suppression. Neuronal activity is inherently noisy ^71–73^, and inappropriate muscle activation could impair locomotor efficiency and reduce fitness. A continuous drive onto inhibitory premotor interneurons may suppress spurious contractions, providing a form of tonic inhibition that is distinct from classical reciprocal inhibition and habituation mechanisms.

A second possible function is the regulation of muscle tone. Even in the absence of contractions, body-wall muscles generate baseline tension that stabilizes posture and supports efficient force production ^74–76^. Modulation of A02l tonic activity may tune this baseline tension to shape axial kinematics during locomotion. These hypotheses warrant further investigation, particularly by quantifying how A02l activity influences passive tension and stiffness in longitudinal muscles, and by testing whether perturbing tonic inhibition via A02l alters force transmission, stride efficiency, or locomotor robustness under conditions of sensory or neural noise.

### Relationship between A02l neurons and PMSIs

The soma of A02l neurons is located in the ventromedial region of the cortex, and their axons project dorsally and then turn along the dorsal side of the neuropil. This morphology resembles that of *period*-positive median segmental interneurons (PMSIs), which derive from neuroblast lineage 2. PMSIs (A02a-j) are glutamatergic interneurons that include first-order premotor neurons directly inhibiting motor neurons and promoting muscle relaxation following segmental contraction during crawling ^77^. Although the neuroblast lineage of A02l remains unclear ^78^, A02l neurons and PMSIs have somata in a similar location and exhibit similar proximal axon trajectories, and both contribute to muscle relaxation, suggesting a degree of functional convergence. Whereas PMSIs act as premotor interneurons that directly inhibit motor neurons, A02l neurons appear to promote relaxation of longitudinal muscles indirectly by activating multiple inhibitory premotor interneurons. Future work dissecting the molecular identities, developmental programs, and circuit functions of these two interneuron classes will clarify how distinct architectures are recruited to achieve muscle relaxation during locomotion.

### Orthogonal regulation of axial locomotion

A notable feature of A02l connectivity is its preferential targeting of longitudinal muscle-innervating premotor circuits, which drive segment shortening and forward propulsion. We previously identified the A31c-A26f circuit as a regulator of transverse muscle contraction, which sets interwave duration and thereby locomotion speed ^33^. Although both circuits use precisely timed release of inhibition to shape kinematic phase relationships, they operate through distinct modes: A02l exhibits tonic activity with transient suppression, whereas the A31c-A26f circuit relies on transient disinhibition.

Because longitudinal and transverse muscles are not contracted simultaneously ^33,44,79^, these two muscle groups can be regarded as functionally antagonistic, analogous to flexor-extensor pairs in vertebrates. Regulation of such antagonistic muscle groups by higher-order promoter interneurons via the A02l pathway may underlie the temporal control of coordinated contraction patterns. Connectomics analysis revealed that A02l neurons receive descending and intersegmental inputs. The descending input may coordinate antagonistic muscle groups and initiate A02l tonic activity, thereby enforcing tonic inhibition of muscle contraction during quiescent periods and potentially reducing energetic costs.

It remains unclear why longitudinal muscles are controlled through tonic inhibition with transient relief, whereas transverse muscles are regulated via temporally gated inhibition. The distinction may reflect differing functional demands, such as a requirement for noise robustness, biomechanical constraints, or distinct modes of descending control.

### Dedicated or general function of A02l neurons

Although we focused on crawling, larvae exhibit a rich repertoire of behaviors, including bending, turning, sweeping, hunching, rearing, and rolling. Given the roles of A02l neurons in the global suppression of muscle contraction, it will be important to determine whether its tonic activity is maintained across behavioral states or instead shows behavior-specific patterns of suppression associated with non-crawling actions. Alternatively, transient suppression of A02l may itself be instructive, directly triggering distinct motor programs. For example, the whole-body contraction induced by acute A02l inhibition (**Figure 6B**) may correspond to a hunching-like behavior.

Future *in vivo* imaging across multiple behavioral contexts (e.g., using the tracking preparation shown in **Figure 1L**) will help distinguish whether A02l functions primarily as a global gating mechanism or plays a more direct role in specifying particular motor patterns. Such analyses will deepen our understanding of how tonic neuronal inhibition contributes to smooth behavioral transitions and action selection in motor control.

## STAR Methods

### Fly strains

Fly larvae were raised at 25 °C unless otherwise noted. The following *Drosophila* melanogaster lines were used in this research.

● *w1118* (Bloomington *Drosophila* Stock Center [BDSC], #3605)
● *R84G07-p65.AD (attP40); R95H02-Gal4.DBD/TM6B, Tb* (A02l-split1; specifically labels A02l neurons in A1 – A7 neuromeres)
● *R13B07-p65.AD (attP40)/CyO, RFP, Tb; R84G07s-Gal4.DBD/TM6B, Tb* (A02l-split2; specifically labels A02l neurons in A4-A7 neuromeres)
● *R95H02-Gal4 (attP2)* (A02l-Gal4, BDSC #40716)
● *eve[RRa-F]-GAL4* (targets aCC/RP2 motor neurons, a gift from Dr. Miki Fujioka, now available as BDSC #7470)
● *UAS-CD4::GCaMP6f (attP40)* (Kyoto Stock Center, #118813)
● *UAS-CD4::GCaMP6f (su(Hw)attP5)* (Kyoto Stock Center, #118814)
● *UAS-GCaMP6f (attP40)* (BDSC #42747)
● *UAS-6x-mCherry (attP2)* (BDSC #52268)
● *UAS-CD4::jRGECO1b (VK5)* (generated in this study; see below)
● *UAS-ASAP5 (attP40)/CyO; UAS-jRGECO1a (VK5)/TM6B, Tb* (BDSC #605338)
● *UAS-iGABASnFR2 (attP40)* (a gift from Dr. Vivek Jayaraman)
● *UAS-GRAB-ACh3.0 (attP40)* (BDSC #86549)
● *UAS-CsChrimson::mVenus (attP18)* (BDSC #55134)
● *UAS > dsFRT > CsChrimson::mVenus (attP18), pBPhsFlp2::Pest (attP3)* (for clonal optogenetic activation; gift from Drs. Stefan Pulver, Karen Hibbard, and the Rubin lab)
● *UAS-GtACR1::eYFP (attP2)* (gift from Dr. Chris Doe, now available as BDSC #92983)
● *MCFO-4* (BDSC #64088)

### Construction of UAS-CD4::jRGECO1b

The coding sequence for a CD4-jRGECO1b fusion protein, consisting of CD4 ^80^ and jRGECO1b ^39^, was codon-optimized for *Drosophila melanogaster* (Bio Basic Inc, Canada). To generate the UAS-CD4::jRGECO1b construct, the CD4::jRGECO1b coding sequence, together with the Syn21 translational enhancer at the 5’ end was cloned into the pJRFC28-10xUAS-IVS-GFP-p10 plasmid backbone ^81^. The resulting transgene was inserted into the VK00005 landing site by PhiC31-mediated integration (BestGene Inc., USA) to generate UAS-CD4::jRGECO1b transgenic fly lines.

### Immunohistochemistry

Third-instar larvae were dissected in phosphate-buffered saline (PBS), and isolated central nervous systems were mounted on MAS-coated slide glass (Matsunami Glass, Japan). Samples were fixed in 4% formaldehyde for 30 min, washed twice in PBS containing 0.2% Triton X-100 (PBT) for 15 min, and blocked in 5% normal goat serum in PBT for 30 min at room temperature. Preparations were incubated with primary antibodies at 4°C overnight, washed twice with PBT for 15 min, and incubated with secondary antibodies at 4°C overnight. After two final washes in PBT for 15 min each, the solution was replaced with PBS. Samples were imaged using a confocal microscope (FV3000, Evident, Japan) with either a 20x water-immersion objective (XLUMPLFLN 20XW, NA=1.0, Evident, Japan) or a 60x water-immersion objective (LUMPLFLN 60XW, NA=1.0, Evident, Japan).

Antibodies and dilutions used are listed below:

● Anti-GFP, rabbit (Frontier Institute, Af2020): 1:1000
● Anti-Fas2, mouse (Developmental Studies Hybridoma Bank (DSHB), 1D4): 1:100
● Anti-V5, mouse (Invitrogen, R960-25): 1:500
● Anti-HRP-Alexa647 (Jackson ImmunoResearch): 1:300
● Anti-HA, rabbit (Cell Signaling Technology, C29F4): 1:1000
● Anti-ChAT, mouse (DSHB, 4B1): 1:50
● Anti-vGluT ^82^, rabbit (gift from Dr. Hermann Aberle): 1:1000
● Anti-vGAT ^83^, rabbit (gift from Dr. David Krantz): 1:1000
● Goat anti-rabbit Alexa Fluor 488 (Invitrogen, A11034): 1:300
● Goat anti-guinea pig Alexa Fluor 488 (Invitrogen, A11073): 1:300
● Goat anti-mouse Alexa Fluor 555 (Invitrogen, A21424): 1:300
● Goat anti-rabbit Cy5 (Invitrogen, A10523): 1:300

### Functional imaging of the isolated central nervous system

Wandering third-instar larvae were dissected in TES buffer (5 mM TES, 135 mM NaCl, 5 mM KCl, 4 mM MgCl_2_, 2 mM CaCl_2_, 36 mM sucrose, pH = 7.15). The isolated central nervous system was mounted on a MAS-coated slide glass (S9215, Matsunami Glass, Japan) and immersed in TES buffer. Fluorescence signals from the VNC were recorded using a spinning disk confocal unit (CSU W1, Yokogawa, Japan) and an EMCCD camera (iXon, Andor Technology, Germany) mounted on an upright microscope (BX51WI, Evident, Japan) equipped with a 20x objective (UMPlanFLN20X, NA=0.5, Olympus, Japan).

For imaging two z-planes, a piezo objective scanner (P-725.2CL, PI, Germany) was used. Single-color calcium imaging with GCaMP was performed using a 488-nm laser (OBIS, Coherent, USA), and dual-color imaging additionally used a 561-nm laser (Votran, USA). Green and red fluorescence emissions were separated using a dual-view system (DV2, Photometrics, USA) mounted on CSU-W1.

Image acquisition rates were 33 frames per second (fps) for single-plane calcium and voltage imaging, 10 fps for acetylcholine imaging and calcium imaging during TTX applications, and 8 fps per plane for two-plane imaging.

For pharmacological experiments, calcium imaging was first performed in TES buffer for 30 s. The external solution was then exchanged for TES buffer supplemented with 2 μM TTX (1069, Tocris Bioscience), followed by a 3-minute incubation. Calcium imaging was subsequently repeated for an additional 30 s.

### Analysis of calcium, neurotransmitter, or voltage imaging from the isolated larval CNS

Regions of interest (ROIs) were selected in the isolated larval CNS, corresponding to A02l presynaptic sites or motor neuron (MN) dendrites in each neuromere, and the mean fluorescence signal *f* was computed for each ROI. Baseline fluorescence *f*_*base*_(*t*) was defined as a low-pass-filtered *f* using a fourth-order Butterworth filter (*f*_*cutoff*_ = *0*.*02* Hz). Normalized fluorescence changes were calculated as *Δf*/*f* = (*f* − *f*_*base*_)/*f*_*base*_. For experiments involving simultaneous calcium imaging of A02l neurons and aCC motor neurons, calcium peaks in motor neurons were detected using the find_peak function in SciPy. Traces from individual animals were temporally interpolated, and averaged A02l *Δf*/*f* waveforms were computed relative to the timing of motor neuron calcium peaks within the same hemisegment. Cross-correlation functions between A02l and motor neuron activity were calculated using the correlate function in scipy.signal package.

In the hyperpolarizing experiments with Kir2.1 and pharmacological experiments with tetrodotoxin (TTX), we defined baseline fluorescence *f_base_* as the average baseline fluorescence (computed by a Butterworth filter) throughout the recording. For the control group in Kir 2.1 experiments or recordings in normal saline, dip fluorescence *f_dip_* was defined as the minimum fluorescence during calcium dips.

### *In vivo* calcium imaging from intact crawling larvae

Calcium recordings were performed as previously described ^37^ using a custom two-photon microscope equipped with dual acousto-optic deflectors (AODs) and a tunable acoustic gradient (TAG) lens for random-access scanning. For *in vivo* recording, mCherry was expressed for tracking and GCaMP6f for neural activity monitoring in A02l neurons using either A02l-split1 or R19C05-Gal4. Larvae were placed on an agar-coated coverslip mounted on a 3-axis stage. To identify and select target neurons, larvae were initially immobilized using a reversible immobilization chamber on the stage. This chamber allowed precise control of the gap between the coverslip and an acrylic window; once neurons were selected, the gap was increased to allow crawling behavior during the experiment.

A Ti:sapphire laser (Chameleon Ultra II, Coherent) tuned to 960 nm was used to excite both GCaMP6f and mCherry. Two adjacent cell bodies or axon terminals were tracked using the mCherry signal. Larval movements were recorded from below using an infrared camera (Basler acA640-90um) under IR illumination (Thorlabs Fiber-Coupled LED, M850F2) to determine the timing and direction of each locomotor bout.

For analysis, fluorescence was extracted from each ROI in the green and red channels, and the ratiometric signal was computed as *R*(*t*) = *f*_*green*_(*t*)/*f*_*red*_(*t*), where green and red fluorescence correspond to GCaMP6f and mCherry signals, respectively. For each recording, the baseline *R_0_* was defined as 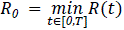, where *T* is the duration of the recording window, and traces were plotted as *R*/*R_0_*. Traces shown in Figures 1 and S2 correspond to these baseline-corrected ratiometric measures.

### Optogenetic behavioral experiments in intact larvae

Larvae were raised at 25 °C. Second- or early third-instar larvae were transferred to yeast paste supplemented with all-trans-retinal (ATR; 3 mM) and reared for 24 – 48 h in darkness. Larvae were rinsed with water and placed on apple juice agar plates for a 2-min acclimation period before recording.

Larval movements were recorded using a CCD camera (XCD V60, Sony, Japan) under infrared illumination (850 nm; LDQ-150IR2-850, CCS, JAPAN) at ∼40 μW/mm^2^. For CsChrimson activation, larvae were exposed to red light (660 nm LED, M660L3, Thorlabs, USA) at 60 μW/mm^2^ for 5 s. For GtACR1-mediated inhibition, green light (250 μW/mm^2^) from a mercury lamp (U-HGLGPS, Evident, Japan) was applied through a 530-550 nm bandpass filter for 5 s.

### Analysis of optogenetics experiments

For CsChrimson activation experiments, forward or backward crawling events were manually counted. The number of crawling events during the 5 s before and after the onset of red-light stimulation was compared.

For GtACR1 inhibition experiments, larval body area was automatically segmented in each frame after Gaussian blurring (σ = 3 pixels) using the threshold_li function implemented in scikit-image ^84^. Body area was normalized to the pre-stimulation baseline, and relative changes were used to quantify whole-body contraction.

### EM connectomics analysis

Electron microscopy (EM) connectomics analysis was performed using the larval central nervous system dataset of a first-instar larva ^26,27^. To examine the synaptic connectivity of A02l neurons, we newly reconstructed and curated portions of the circuit in this study, focusing on both upstream (presynaptic) and downstream (postsynaptic) partners of A02l neurons in the A1 neuromere. Synaptic connections were identified based on established ultrastructural criteria ^26^. To characterize the functional output of the A02l circuit, we further analyzed connections from A02l neurons to motor neurons and from motor neurons to muscles. Motor neuron identities and their target muscles were assigned based on an established motor neuron–muscle map ^52^. This allows us to infer which muscles are indirectly inhibited by A02l neurons through their downstream circuitry. The circuit diagram in **Figure S6F** was generated using a custom Python code.

### Statistical tests

Statistical analyses were performed in Python using Scipy (scipy.stats) and Statsmodels (statsmodels.stats.multitest). Statistical significance is denoted as *** p < 0.001, ** p < 0.01, * p < 0.05, and n.s., not significant.

## Resource Availability

### Lead contact

Requests for further information and resources should be directed to and will be fulfilled by the lead contact, Hiroshi Kohsaka.

### Materials availability

Materials that are not publicly available are available upon request to the lead contact.

### Data and code availability

The data and codes are available through https://github.com/takahisad/Date_2026_A02l.

Any additional information required to analyze the data reported in this paper is available from the lead contact upon request.

## Acknowledgment

We thank Drs. Chris Doe, Miki Fujioka, James Truman, Vivek Jayaraman for the fly stocks. We thank Dr. Albert Cardona for providing access to the 1st instar larval EM dataset. We thank Drs. Hermann Aberle and David Krantz for the antibodies. T.D. was supported by Forefront Physics and Mathematics Program to Drive Transformation (FoPM), a World-leading Innovative Graduate Study (WINGS) Program, the University of Tokyo. This work was supported by MEXT/JSPS KAKENHI grants (18H05113, 19H04742, 20H05048, 21H05675, 21H02576, 22K19479, 22H05487, 23H04213, 24H01225, 24K02117, 25H02486 to A.N.; 17K07042, 20K06908, 21H05301, 23K05959 to H.K.), by the BBSRC (BB/W018675/1 to M.F.Z.), by NIH (1DP2EB022359-01 to M.G.), and by NSF (1455015 to M. G.).

## Author Contribution

Conceptualization, T.D., A.N., M.F.Z., and H.K.; data curation, T.D., M.F.Z., and H.K., formal analysis, T.D., M.F.Z., and H.K., funding acquisition, M.G., A.N., M.F.Z., and H.K.; investigation, T.D., Y.L., A.Y., P.M., R.W., M.G., M.F.Z., and H.K.; project administration, H.K.; resources, A.N. and H.K., supervision, M.F.Z. and H.K.; visualization, T.D., M.F.Z., and H.K.; writing – original draft, T.D., M.F.Z., and H.K.; writing – review & editing, T.D., Y.L., A.Y., M.G., M.F.Z., and H.K.

## Supplementary information

Figures S1-S7

Tables S1 and S2

Movies S1-S5

---

**Figure S1:**
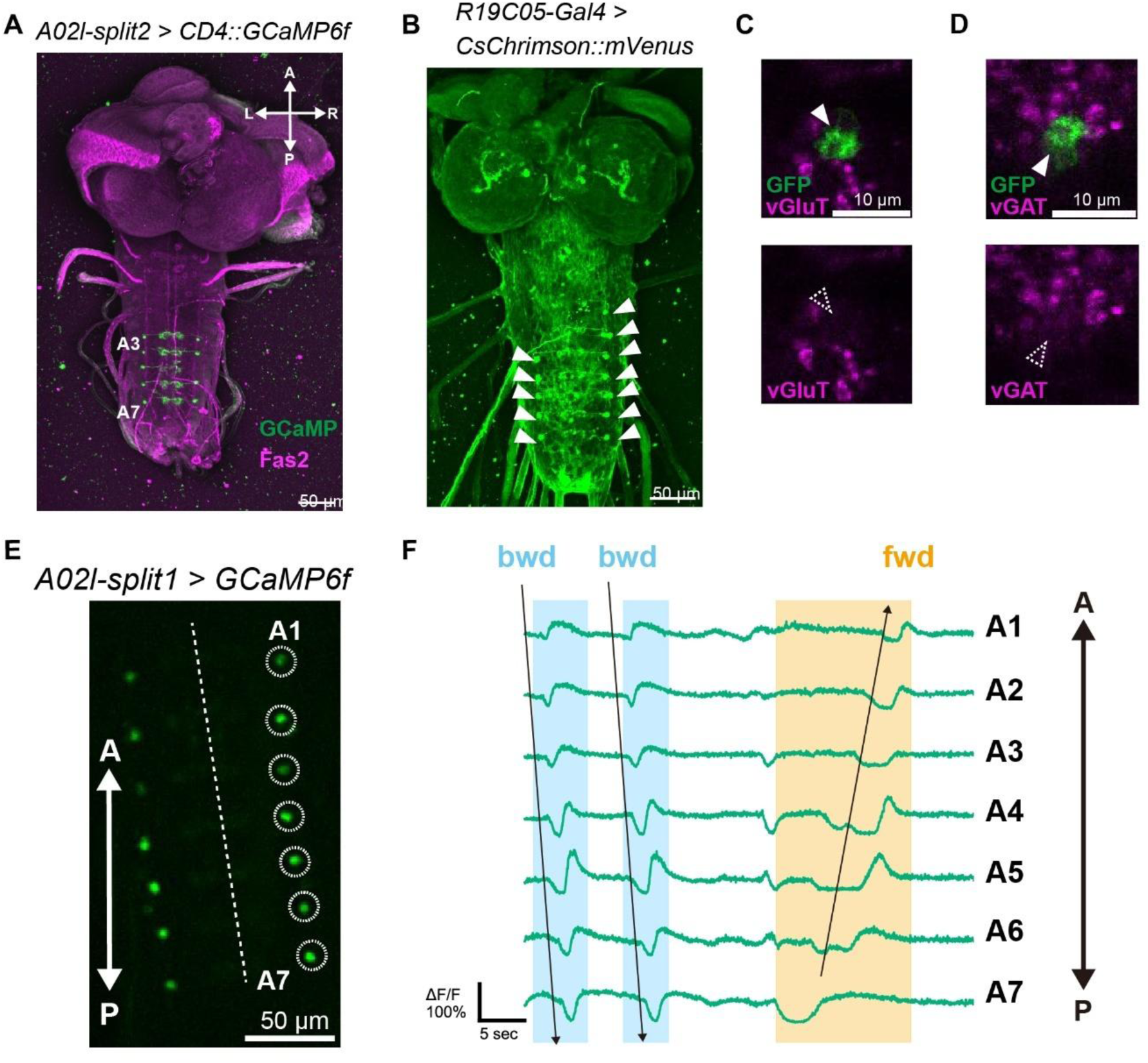
Expression patterns of A02l-Gal4 lines and immunostaining of A02l neurons for glutamate and GABA markers. **(A)** Expression pattern of A02l-split2 (*R13B07-AD; R84G07-DBD > CD4::GCaMP6f*), labeling A02l neurons in A3-A7 neuromeres. A, anterior; P, posterior; L, left; R, right. **(B)** Expression pattern of R19C05-Gal4 (*R19C05-Gal4 > CsChrimson::mVenus*). **(C)** A02l axon terminals (arrowhead; *A02l-split1 > CD4::GCaMP6f*) are not co-labeled with vesicular glutamate transporter (vGluT). Top, merged image; bottom, vGluT channel alone. **(D)** A02l axon terminals (arrowhead, *A02l-split1 > CD4::GCaMP6f*) are not co-labeled with vesicular GABA transporter (vGAT). Top, merged image; bottom, vGAT channel alone. **(E)** Representative calcium imaging frame from *A02l-split1 > CD4::GCaMP6f* larvae. GCaMP signals were extracted from A02l axon terminals in the A1-A7 neuromeres (dashed circles). The dashed line indicates the VNC midline. A, anterior; P, posterior. **(F)** Representative calcium traces from A02l axon terminals in the A1-7 neuromeres (*A02l-split1 > CD4::GCaMP6f*, the larva in panel E) during fictive locomotion. Two backward (bwd) waves and one forward (fwd) wave are indicated by arrows and shaded bands. A, anterior; P, posterior.

**Figure S2.**
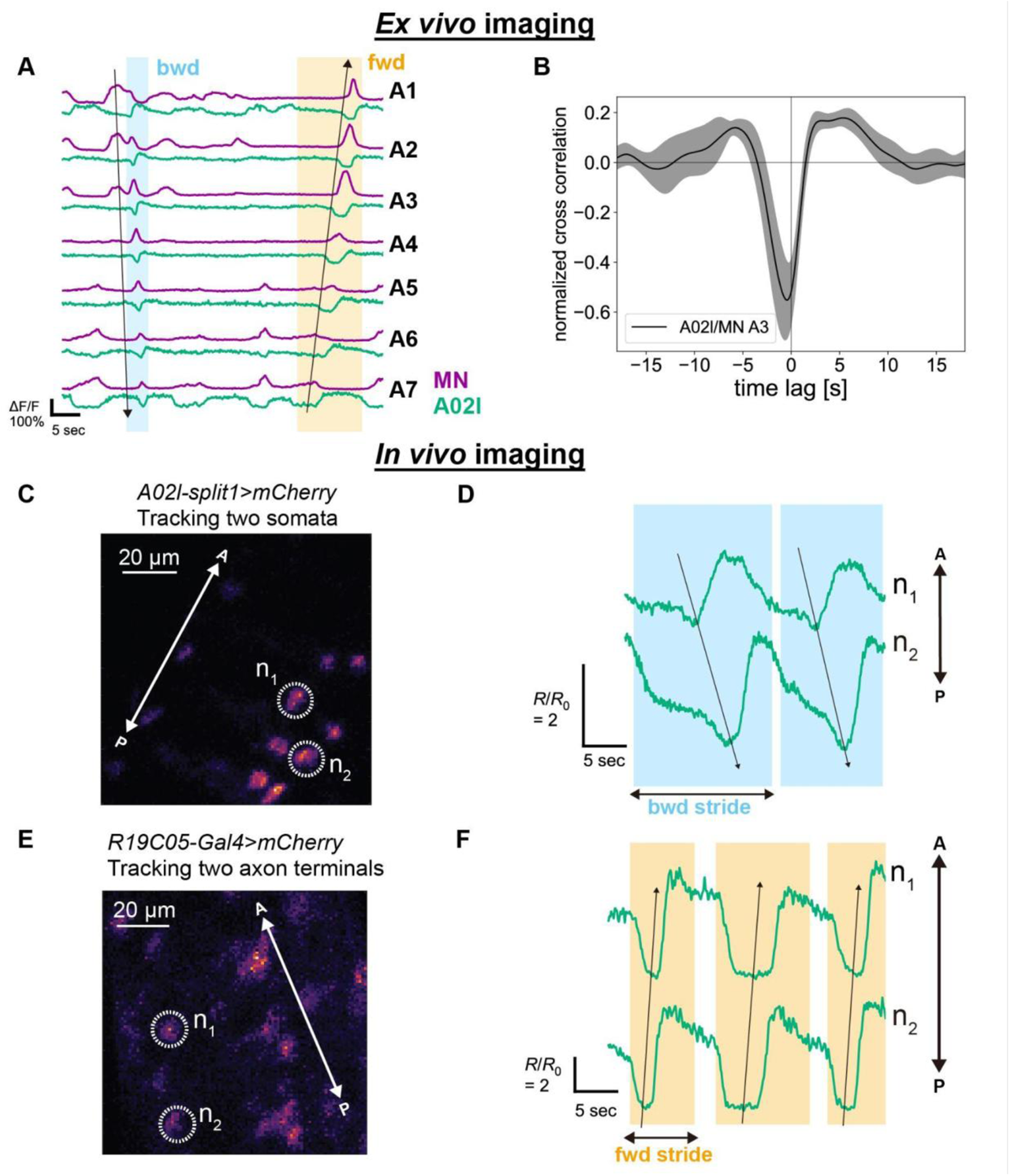
Ex vivo and in vivo calcium imaging of A02l neurons. **(A)** Representative calcium traces of aCC motor neurons (magenta) and A02l neurons (green) in the A1 –A7 neuromeres. Forward (fwd) and backward (bwd) fictive locomotor waves are indicated by arrows and shaded bands. A02l traces show calcium dips coincident with motor neuron activation in the corresponding segment. **(B)** Normalized cross-correlation of Ca^2+^ activity between A02l neurons and motor neurons in segment A3 (16 A02l/MN pairs from 8 animals). The trough at zero lag (*t_lag_* = *0*) indicates anticorrelation between A02l and motor neuron activity. (**C, D**) Representative *in vivo* calcium imaging of A02l neurons using A02l-split1 during backward locomotion. (**C)** Image of two tracked A02l somata; n1 is located in the neuromere immediately anterior to n2. (**D)** Corresponding calcium traces from the neurons shown in (C) (*R* = *f_GCaMP_*/*f_mCherry_*, normalized to the minimum value *R_0_*). Calcium dips propagate posteriorly during backward crawling. A, anterior; P, posterior. (**E, F**) Representative *in vivo* calcium imaging of A02l neurons using R19C05-Gal4 during forward locomotion. (**E)** Image of two tracked A02l axon terminals; n1 is located in the neuromere immediately anterior to n2. (**F)**: Corresponding calcium traces from the neurons shown in (E) (*R* = *f_GCaMP_*/*f_mCherry_*, normalized to the minimum value *R_0_*). Calcium dips propagate anteriorly during forward crawling. A, anterior; P, posterior.

**Figure S3.**
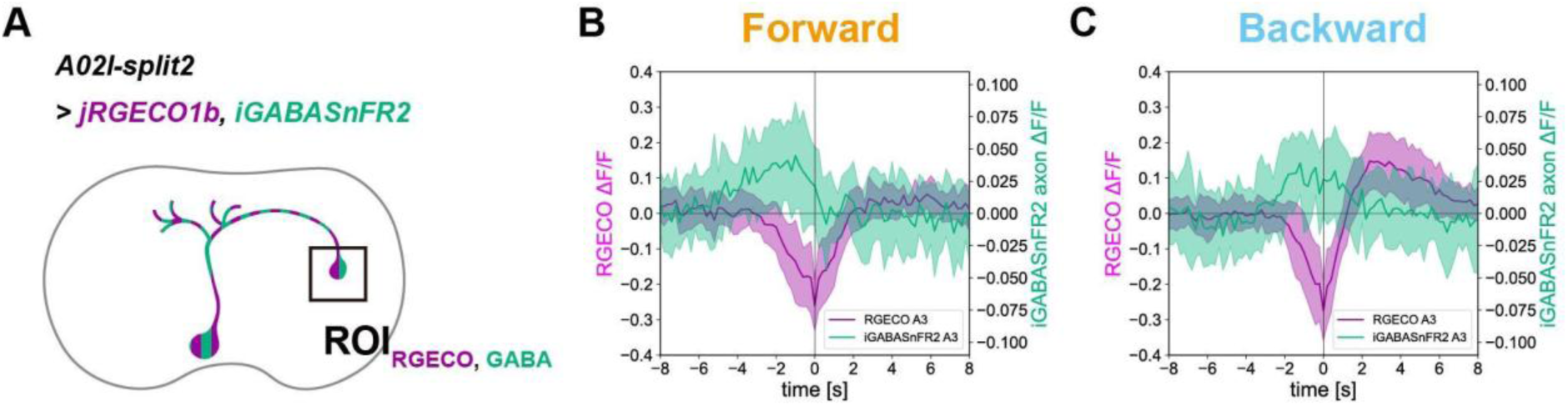
GABA signals at A02l axon terminals. **(A)** Schematic indicating the ROIs for simultaneous calcium and GABA imaging at A02l axon terminals (*A02l-split2 > jRGECO1b, iGABASnFR2*). Both jRGECO1b and iGABASnFR2 signals were analyzed in axon terminals. **(B, C)** Averaged ΔF/F traces of jRGECO1b (magenta) and iGABASnFR2 (green) from A02l axon terminals in the A3 neuromere during forward waves (**B**, 17 events from 4 animals) and backward waves (**C**, 18 events from 6 animals). Time 0 corresponds to the negative peak of A02l activity. Shaded bands indicate standard deviation.

**Figure S4.**
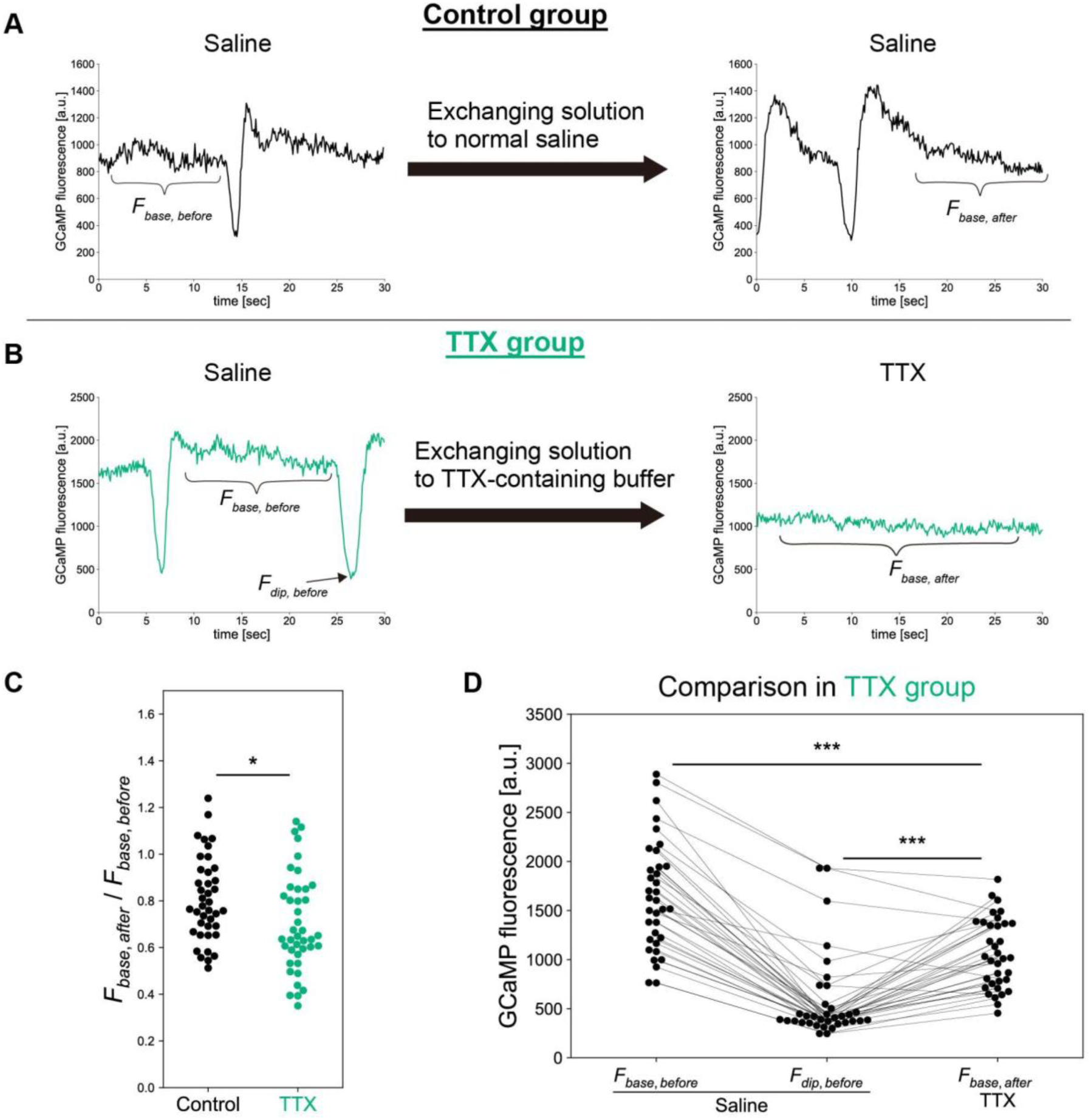
Tetrodotoxin moderately reduces tonic calcium levels in A02l neurons. **(A)** Representative traces of the control group. Genotype: *A02l-split1 > CD4::GCaMP6f.* Left: representative calcium trace from an A02l axon terminal in the A4 neuromere during *ex vivo* imaging in normal saline. Baseline fluorescence (*f_base,before_*) was defined as the mean fluorescence during rest. Right: calcium trace from the same axon terminal three minutes after exchange to the same normal saline. Locomotor activity was maintained, and the A02l neuron showed calcium dips. Post-buffer-exchange baseline fluorescence (*f_base,after_*) was defined as the mean fluorescence during rest. **(B)** Representative traces of the experimental group supplemented with tetrodotoxin (TTX). Genotype: *A02l-split1 > CD4::GCaMP6f.* Left: representative calcium trace from an A02l axon terminal in the A4 neuromere during *ex vivo* imaging in normal saline. Baseline fluorescence (*f_base,before_*) was defined as the mean fluorescence during rest, and dip fluorescence (*f_dip,before_*) as the minimum fluorescence during a calcium dip. Right: calcium trace from the same axon terminal three minutes after exchange to saline containing 2 μM tetrodotoxin (TTX). Locomotor activity was abolished, and no calcium dips were observed. Post-TTX baseline fluorescence (*f_base,after_*) was defined as the mean fluorescence. **(C)** Comparison of GCaMP baseline fold changes (*f_base,after_*/ *f_base,before_*) between control neurons maintained in normal saline (n = 40 cells from 5 animals) and neurons exposed to TTX (n=41 cells from 5 animals). Each point represents a single A02l neuron. TTX significantly reduced baseline GCaMP fluorescence. Mann-Whitney U-test. * p < 0.05. **(D)** Comparison of *f_base,before_*, *f_dip,before_*, and *f_base,after_* within the TTX group (n=35 cells from 5 animals). Values from the same neuron are connected by lines. Although baseline fluorescence decreased after TTX application, the post-TTX baselines *f_base,after_* remained higher than *f_dip,before_*, indicating that tonic A02l activity was not completely abolished. Wilcoxon signed-rank test followed by Holm-Bonferroni correction. *** p < 0.001.

**Figure S5:**
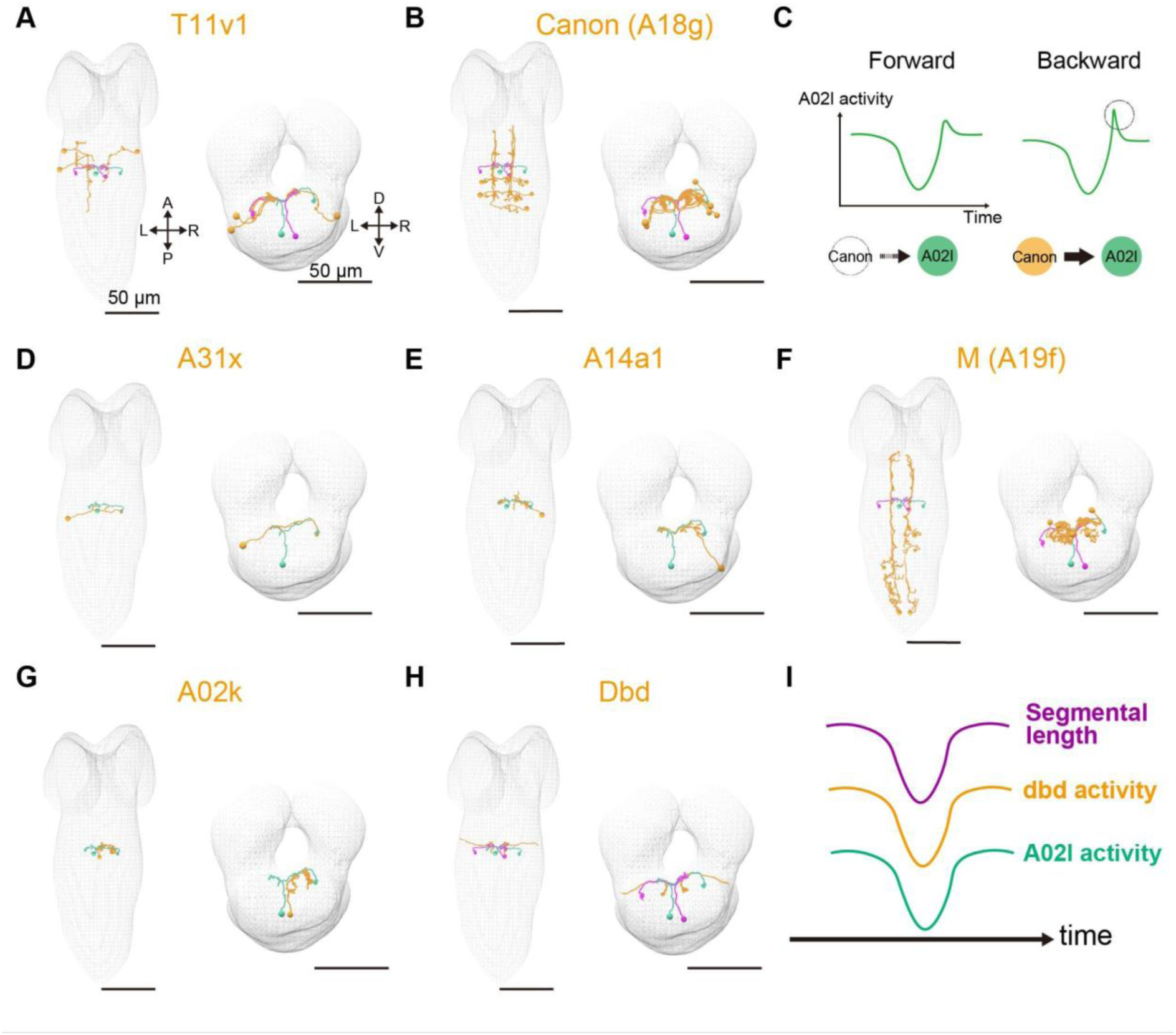
Presynaptic partners of A02l neurons. Morphology of presynaptic partners of A02l neurons reconstructed from EM data (orange). A02l neurons in the left A1 neuromere are shown in green; those in the right A1 neuromere are shown in magenta. Left, dorsal view; right, frontal view. A, anterior; P, posterior; D, dorsal; V, ventral; L, left; R, right. **(A)** T11v1 neurons in the T2-T3 neuromeres. **(B)** Canon (A18g) neurons in the A2-A4 neuromeres. **(C)** Schematic illustrating how Canon neuron activity may contribute to the larger rebound of A02l activity during backward waves. **(D)** A31x neurons in the left A1 neuromere. **(E)** A14a1 neurons in the right A1 neuromere. **(F)** M (A19f) neurons in the A4-A8 neuromeres. **(G)** A02k neurons in the right A1 neuromere. **(H)** dbd neurons in the A1 neuromere. **(I)** Activity patterns of dbd neurons ^49^. Magenta, a segmental length; blue, dbd neuron activity; green, A02l activity. dbd neurons are transiently inhibited near the peak of muscle contraction, coinciding with the calcium dip of A02l. Scale bars, 50 μm.

**Figure S6:**
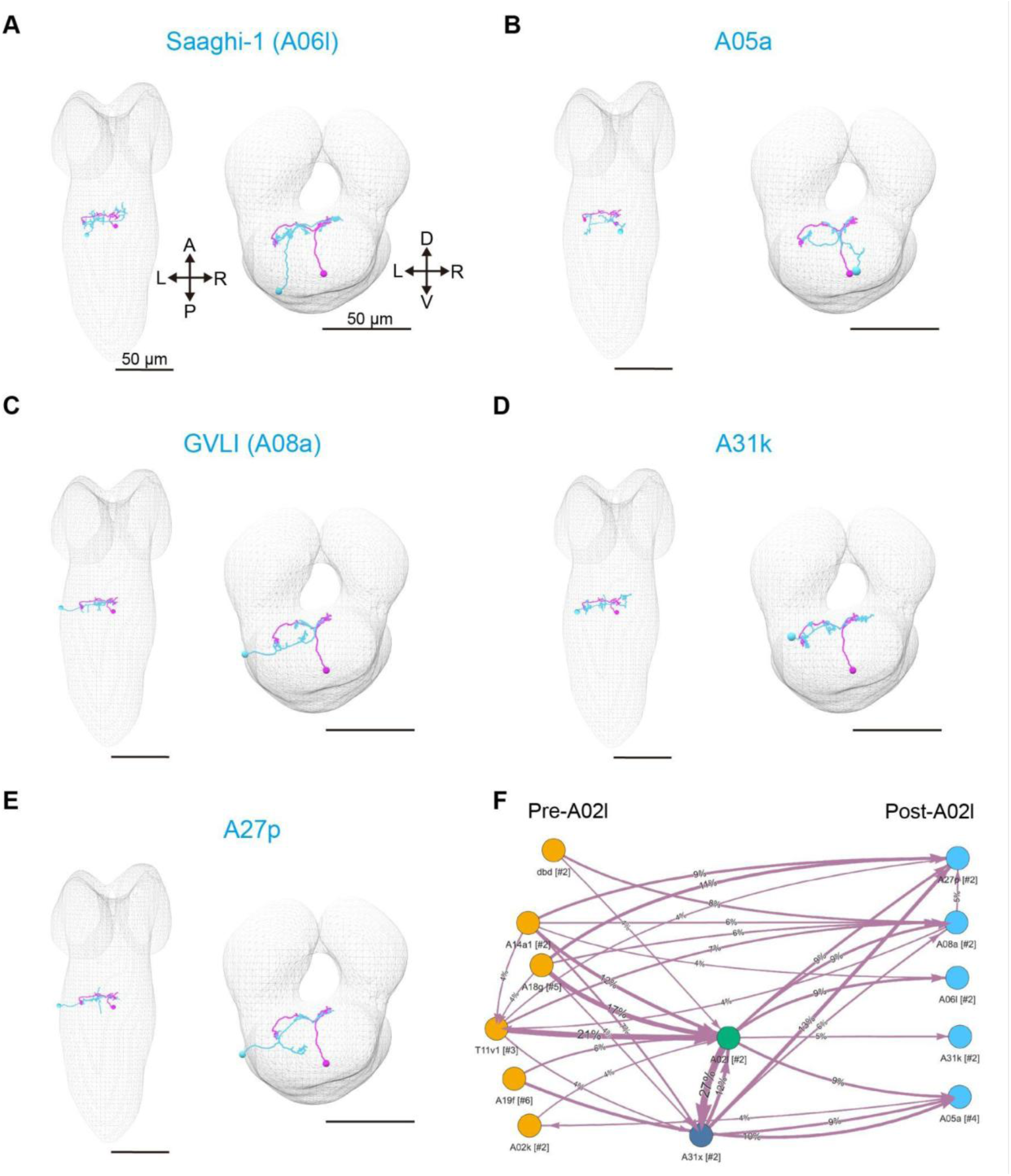
Postsynaptic partners of A02l neurons. Morphology of postsynaptic partners of A02l neurons reconstructed from EM data (cyan). The A02l neuron in the right A1 neuromere is shown in magenta. Left, dorsal view; right, frontal view. A, anterior; P, posterior; D, dorsal; V, ventral; L, left; R, right. **(A)** Saaghi-1 (A06l) neuron in the left A1 neuromere. **(B)** A05a neuron in the right A1 neuromere. **(C)** GVLI (A08a) neuron in the left A1 neuromere. **(D)** A31k neuron in the left A1 neuromere. **(E)** A27p neuron in the left A1 neuromere. Scale bars, 50 μm. **(F)** Circuit diagram of A02l neurons in the A1 neuromere and their postsynaptic partners. Each node represents a single cell type, and “[#…]” indicates the number of neurons included in each node. The weight of each edge is the synapse occupation rate of the postsynaptic neuron. Connections with lower than 3.5% synapse occupation rates were omitted for clarity.

**Figure S7.**
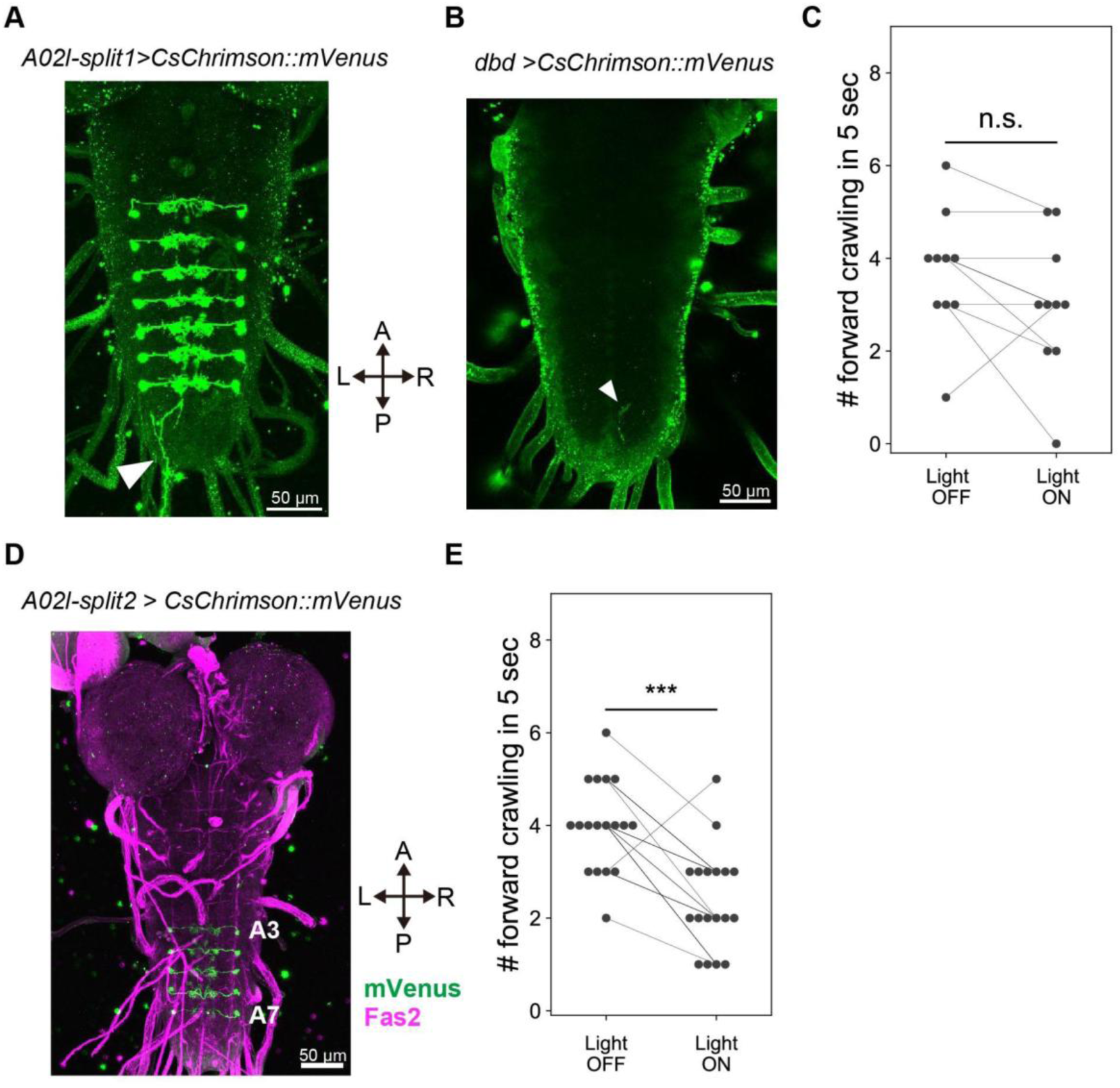
Optogenetic activation of A02l neurons decreases the number of crawling events. **(A)** Expression pattern of *A02l-split1 > CsChrimson::mVenus (attP2)*. A02l-split1 stochastically labels several dbd sensory neurons (white arrowhead). A, anterior; P, posterior; L, left; R, right. **(B)** A dbd neuron stochastically labeled using the clonal optogenetic activation method (white arrowhead). Genotype: *A02l-split1, UAS > dsFRT > CsChrimson::mVenus; pBPhsFlp2::Pest.* Animals were fed ATR. **(C)** Comparison of the number of crawls during the 5 s before and after photoactivation in animals with CsChrimson expression in dbd sensory neurons (n = 10 animals). Wilcoxon signed-rank test; n.s., p > 0.05. **(D)** Expression pattern of *A02l-split2 > CsChrimson::mVenus (attP18)*. A02l-split2 stochastically labels A3 –A7 A02l neurons. A, anterior; P, posterior; L, left; R, right. **(E)** Comparison of the number of crawls during the 5 s before and after photoactivation in *A02l-split2 > CsChrimson::mVenus (attP18)* (n = 18 animals). Animals fed ATR. Wilcoxon signed-rank test. *** p < 0.001.

**Supplementary Table 1.** Presynaptic neurons of A02l neurons in the A1 neuromere identified by EM reconstruction. This table lists 24 upstream neurons that form two or more synaptic connections with A02l neurons in the A1 neuromere. Columns, from left to right, indicate the presynaptic neuron name, the total number of synaptic connections with A02l neurons in the A1 neuromere, and the number of connections onto the left-side A02l neuron (A02l_a1l) and right-side A02l neuron (A02l_a1r), respectively.

| Upstream neuron | Sum | #Synapses with A02l_a1l | #Synapses with A02l_a1r |
| --- | --- | --- | --- |
| T11v1_t2r | 13 | 4 | 9 |
| A14a1_a1l | 8 | 0 | 8 |
| A31x_a1l | 8 | 8 | 0 |
| A18g_a3l | 8 | 3 | 5 |
| A18g_a2l | 8 | 2 | 6 |
| A14a1_a1r | 7 | 7 | 0 |
| A31x_a1r | 7 | 0 | 7 |
| T11v1_t3l | 7 | 5 | 2 |
| T11v1_t2l | 6 | 6 | 0 |
| A14?_a2l | 4 | 0 | 4 |
| A02k_a1l | 3 | 2 | 1 |
| A18g_a2r | 3 | 3 | 0 |
| dbd_a1l | 3 | 2 | 1 |
| A01c1_a1r | 2 | 0 | 2 |
| A03a6c?_a1r | 2 | 0 | 2 |
| dbd_a1r | 2 | 0 | 2 |
| A19f_a6l | 2 | 2 | 0 |
| a11x_t3l | 2 | 2 | 0 |
| A02k_a1r | 2 | 0 | 2 |
| A31f_a1r? | 2 | 0 | 2 |
| A03x_a1r | 2 | 1 | 1 |
| A18g_a3r | 2 | 2 | 0 |
| A19f_a8l | 2 | 2 | 0 |
| A08a_a1r | 2 | 0 | 2 |

**Supplementary Table 2.** Postsynaptic neurons of A02l neurons in the A1 neuromere identified by EM reconstruction. This table lists the 27 downstream neurons that form two or more synaptic connections with A02l neurons in the A1 neuromere. Columns, from left to right, indicate the postsynaptic neuron name, the total number of synaptic connections with A02l neurons in the A1 neuromere, and the number of connections from the left-side A02l neuron (A02l_a1l) and right-side A02l neuron (A02l_a1r), respectively.

| Downstream neuron | Sum | #Synapses with A02l_a1l | #Synapses with A02l_a1r |
| --- | --- | --- | --- |
| A06l_a1r (Saaghi-1) | 28 | 28 | 0 |
| A31k_a1r | 21 | 21 | 0 |
| A06l_a1l (Saaghi-1) | 20 | 0 | 20 |
| A08a_a1r | 19 | 19 | 0 |
| A05a_a1r | 16 | 0 | 16 |
| A27p_a1l | 14 | 0 | 14 |
| A08a_a1l | 13 | 0 | 13 |
| A31x_a1r | 12 | 0 | 12 |
| A31x_a1l | 12 | 12 | 0 |
| A31k_a1l | 10 | 0 | 10 |
| A05a_t3l | 9 | 9 | 0 |
| A27p_a1r | 8 | 8 | 0 |
| A05a_t3r | 8 | 0 | 8 |
| A05a_a1l | 8 | 8 | 0 |
| A27k_a1r | 6 | 6 | 0 |
| A02j_a1l | 5 | 0 | 5 |
| A27l_a1l | 4 | 0 | 4 |
| A31f_a1r? | 4 | 0 | 4 |
| A31d_a1l | 4 | 0 | 4 |
| A03x_a1l | 4 | 0 | 4 |
| A02j_a2l | 3 | 0 | 3 |
| A02g_a1r | 3 | 3 | 0 |
| A07x1_a1l | 2 | 0 | 2 |
| A27l_a1r | 2 | 2 | 0 |
| A31d_a1r | 2 | 2 | 0 |
| A31f_a1l? | 2 | 2 | 0 |
| MN1SNb/d-ls RP5_a1l | 2 | 0 | 2 |

## References

1. Goulding, M. Circuits controlling vertebrate locomotion: moving in a new direction. Nat. Rev. Neurosci. 10, 507–518 (2009).

2. Goulding, M., Bourane, S., Garcia-Campmany, L., Dalet, A. & Koch, S. Inhibition downunder: an update from the spinal cord. Curr. Opin. Neurobiol. 26, 161–166 (2014).

3. Talpalar, A. E. et al. Dual-mode operation of neuronal networks involved in left-right alternation. Nature 500, 85–88 (2013).

4. Kiehn, O. Decoding the organization of spinal circuits that control locomotion. Nat. Rev. Neurosci. 17, 224–238 (2016).

5. Johnson, M. D., Hyngstrom, A. S., Manuel, M. & Heckman, C. J. Push-pull control of motor output. J. Neurosci. 32, 4592–4599 (2012).

6. Benjamin, P. R., Staras, K. & Kemenes, G. What roles do tonic inhibition and disinhibition play in the control of motor programs? Front. Behav. Neurosci. 4, 30 (2010).

7. Hikosaka, O. GABAergic output of the basal ganglia. Prog. Brain Res. 160, 209–226 (2007).

8. Grillner, S. & Robertson, B. The basal ganglia over 500 million years. Curr. Biol. 26, R1088–R1100 (2016).

9. Falgairolle, M., Ceccato, J.-C., Seze, M. de, Herbin, M. & Cazalets, J.-R. Metachronal propagation of motor activity. Front. Biosci. (Landmark Ed.) 18, 820–837 (2013).

10. Dubuc, R., Cabelguen, J.-M. & Ryczko, D. Locomotor pattern generation and descending control: a historical perspective. J. Neurophysiol. 130, 401–416 (2023).

11. Wagenaar, D. A. A classic model animal in the 21st century: recent lessons from the leech nervous system. J. Exp. Biol. 218, 3353–3359 (2015).

12. Zhen, M. & Samuel, A. D. T. C. elegans locomotion: small circuits, complex functions. Curr. Opin. Neurobiol. 33, 117–126 (2015).

13. Clark, M. Q., Zarin, A. A., Carreira-Rosario, A. & Doe, C. Q. Neural circuits driving larval locomotion in Drosophila. Neural Dev. 13, 6 (2018).

14. Kohsaka, H. Linking neural circuits to the mechanics of animal behavior in Drosophila larval locomotion. Front. Neural Circuits 17, 1175899 (2023).

15. Berg, E. M., Björnfors, E. R., Pallucchi, I., Picton, L. D. & El Manira, A. Principles governing locomotion in vertebrates: Lessons from zebrafish. Front. Neural Circuits 12, 73 (2018).

16. Roberts, A., Li, W.-C. & Soffe, S. R. How neurons generate behavior in a hatchling amphibian tadpole: an outline. Front. Behav. Neurosci. 4, 16 (2010).

17. Meng, J. et al. A tonically active master neuron modulates mutually exclusive motor states at two timescales. Sci. Adv. 10, eadk0002 (2024).

18. Zagoraiou, L. et al. A cluster of cholinergic premotor interneurons modulates mouse locomotor activity. Neuron 64, 645–662 (2009).

19. Gradwell, M. A. et al. Multimodal sensory control of motor performance by glycinergic interneurons of the mouse spinal cord deep dorsal horn. Neuron 112, 1302–1327.e13 (2024).

20. Zhong, G., Sharma, K. & Harris-Warrick, R. M. Frequency-dependent recruitment of V2a interneurons during fictive locomotion in the mouse spinal cord. Nat. Commun. 2, 274 (2011).

21. Böhm, U. L. et al. Voltage imaging identifies spinal circuits that modulate locomotor adaptation in zebrafish. Neuron 110, 1211–1222.e4 (2022).

22. Pulver, S. R. et al. Imaging fictive locomotor patterns in larval Drosophila. J. Neurophysiol. 114, 2564–2577 (2015).

23. Jenett, A. et al. A GAL4-driver line resource for Drosophila neurobiology. Cell Rep. 2, 991–1001 (2012).

24. Li, H.-H. et al. A GAL4 driver resource for developmental and behavioral studies on the larval CNS of Drosophila. Cell Rep. 8, 897–908 (2014).

25. Meissner, G. W. et al. A split-GAL4 driver line resource for Drosophila CNS cell types. (2024) doi:10.7554/elife.98405.1.

26. Ohyama, T. et al. A multilevel multimodal circuit enhances action selection in Drosophila. Nature 520, 633–639 (2015).

27. Schneider-Mizell, C. M. et al. Quantitative neuroanatomy for connectomics in Drosophila. Elife 5, (2016).

28. Kohsaka, H., Guertin, P. A. & Nose, A. Neural circuits underlying fly larval locomotion. Curr. Pharm. Des. 23, 1722–1733 (2017).

29. Gowda, S. B. M., Salim, S. & Mohammad, F. Anatomy and neural pathways modulating distinct locomotor behaviors in Drosophila larva. Biology (Basel) 10, (2021).

30. Fushiki, A. et al. A circuit mechanism for the propagation of waves of muscle contraction in Drosophila. Elife 5, (2016).

31. Kohsaka, H. et al. Regulation of forward and backward locomotion through intersegmental feedback circuits in Drosophila larvae. Nat. Commun. 10, 2654 (2019).

32. Chen, T.-W. et al. Ultrasensitive fluorescent proteins for imaging neuronal activity. Nature 499, 295–300 (2013).

33. Liu, Y., Hasegawa, E., Nose, A., Zwart, M. F. & Kohsaka, H. Synchronous multi-segmental activity between metachronal waves controls locomotion speed in Drosophila larvae. Elife 12, e83328 (2023).

34. Sales, E. C., Heckman, E. L., Warren, T. L. & Doe, C. Q. Regulation of subcellular dendritic synapse specificity by axon guidance cues. Elife 8, e43478 (2019).

35. Heckman, E. L. & Doe, C. Q. Presynaptic contact and activity opposingly regulate postsynaptic dendrite outgrowth. Elife 11, e82093 (2022).

36. Lemon, W. et al. Whole-central nervous system functional imaging in larval Drosophila. Nat. Commun. 6, 7924 (2015).

37. Yamaguchi, A. et al. Multi-neuronal recording in unrestrained animals with all acousto-optic random-access line-scanning two-photon microscopy. Front. Neurosci. 17, (2023).

38. Nern, A., Pfeiffer, B. D. & Rubin, G. M. Optimized tools for multicolor stochastic labeling reveal diverse stereotyped cell arrangements in the fly visual system. Proc. Natl. Acad. Sci. U. S. A. 112, E2967–76 (2015).

39. Dana, H. et al. Sensitive red protein calcium indicators for imaging neural activity. Elife 5, e12727 (2016).

40. Hao, Y. A. et al. A fast and responsive voltage indicator with enhanced sensitivity for unitary synaptic events. Neuron 112, 3680–3696.e8 (2024).

41. Baines, R. A., Uhler, J. P., Thompson, A., Sweeney, S. T. & Bate, M. Altered electrical properties in Drosophila neurons developing without synaptic transmission. J. Neurosci. 21, 1523–1531 (2001).

42. Kolb, I., et al. iGABASnFR2: Improved genetically encoded protein sensors of GABA. *bioRxiv* (2025) doi:10.1101/2025.03.25.644953.

43. Jing, M. et al. An optimized acetylcholine sensor for monitoring in vivo cholinergic activity. Nat. Methods 17, 1139–1146 (2020).

44. Zwart, M. F. et al. Selective inhibition mediates the sequential recruitment of motor pools. Neuron 91, 615–628 (2016).

45. Hiramoto, A. et al. Regulation of coordinated muscular relaxation in Drosophila larvae by a pattern-regulating intersegmental circuit. Nat. Commun. 12, 2943 (2021).

46. Zeng, X. et al. An electrically coupled pioneer circuit enables motor development via proprioceptive feedback in Drosophila embryos. Curr. Biol. 31, 5327–5340.e5 (2021).

47. Jonaitis, J., et al. Steering from the rear: Coordination of central pattern generators underlying navigation by ascending interneurons. *bioRxiv* (2024) doi:10.1101/2024.06.17.598162.

48. Suslak, T. J. et al. Piezo is essential for amiloride-sensitive stretch-activated mechanotransduction in larval Drosophila dorsal bipolar dendritic sensory neurons. PLoS One 10, e0130969 (2015).

49. Vaadia, R. D. et al. Characterization of proprioceptive system dynamics in behaving Drosophila larvae using high-speed volumetric microscopy. Curr. Biol. 29, 935–944.e4 (2019).

50. Heckscher, E. S. et al. Even-skipped(+) interneurons are core components of a sensorimotor circuit that maintains left-right symmetric muscle contraction amplitude. Neuron 88, 314–329 (2015).

51. Itakura, Y. et al. Identification of inhibitory premotor interneurons activated at a late phase in a motor cycle during Drosophila larval locomotion. PLoS One 10, e0136660 (2015).

52. Zarin, A. A., Mark, B., Cardona, A., Litwin-Kumar, A. & Doe, C. Q. A multilayer circuit architecture for the generation of distinct locomotor behaviors in Drosophila. Elife 8, e51781 (2019).

53. Klapoetke, N. C. et al. Independent optical excitation of distinct neural populations. Nat. Methods 11, 338–346 (2014).

54. Govorunova, E. G., Sineshchekov, O. A., Janz, R., Liu, X. & Spudich, J. L. NEUROSCIENCE. Natural light-gated anion channels: A family of microbial rhodopsins for advanced optogenetics. Science 349, 647–650 (2015).

55. Mohammad, F. et al. Optogenetic inhibition of behavior with anion channelrhodopsins. Nat. Methods 14, 271–274 (2017).

56. Marder, E. & Bucher, D. Central pattern generators and the control of rhythmic movements. Curr. Biol. 11, R986–96 (2001).

57. McCrea, D. A. & Rybak, I. A. Modeling the mammalian locomotor CPG: insights from mistakes and perturbations. Prog. Brain Res. 165, 235–253 (2007).

58. Grillner, S. & Kozlov, A. The CPGs for limbed locomotion-facts and fiction. Int. J. Mol. Sci. 22, 5882 (2021).

59. Attwell, D. & Laughlin, S. B. An energy budget for signaling in the grey matter of the brain. J. Cereb. Blood Flow Metab. 21, 1133–1145 (2001).

60. Howarth, C., Gleeson, P. & Attwell, D. Updated energy budgets for neural computation in the neocortex and cerebellum. J. Cereb. Blood Flow Metab. 32, 1222–1232 (2012).

61. Harris, J. J., Jolivet, R. & Attwell, D. Synaptic energy use and supply. Neuron 75, 762–777 (2012).

62. Rangaraju, V., Calloway, N. & Ryan, T. A. Activity-driven local ATP synthesis is required for synaptic function. Cell 156, 825–835 (2014).

63. Pulido, C. & Ryan, T. A. Synaptic vesicle pools are a major hidden resting metabolic burden of nerve terminals. Sci. Adv. 7, eabi9027 (2021).

64. Müller, P., Draguhn, A. & Egorov, A. V. Persistent sodium currents in neurons: potential mechanisms and pharmacological blockers. Pflugers Arch. 476, 1445–1473 (2024).

65. Simms, B. A. & Zamponi, G. W. Neuronal voltage-gated calcium channels: structure, function, and dysfunction. Neuron 82, 24–45 (2014).

66. Monteil, A. et al. New insights into the physiology and pathophysiology of the atypical sodium leak channel NALCN. Physiol. Rev. 104, 399–472 (2024).

67. Arnsten, A. F. T., Wang, M. J. & Paspalas, C. D. Neuromodulation of thought: flexibilities and vulnerabilities in prefrontal cortical network synapses. Neuron 76, 223–239 (2012).

68. Wilson, J. A. & Phillips, C. E. Locust local nonspiking interneurons which tonically drive antagonistic motor neurons: physiology, morphology, and ultrastructure. J. Comp. Neurol. 204, 21–31 (1982).

69. Siegler, M. V. S. Local interneurones and local interactions in arthropods. J. Exp. Biol. 112, 253–281 (1984).

70. Laurent, P. et al. Decoding a neural circuit controlling global animal state in C. elegans. Elife 4, (2015).

71. Faisal, A. A., Selen, L. P. J. & Wolpert, D. M. Noise in the nervous system. Nat. Rev. Neurosci. 9, 292–303 (2008).

72. Stein, R. B., Gossen, E. R. & Jones, K. E. Neuronal variability: noise or part of the signal? Nat. Rev. Neurosci. 6, 389–397 (2005).

73. Yarom, Y. & Hounsgaard, J. Voltage fluctuations in neurons: signal or noise? Physiol. Rev. 91, 917–929 (2011).

74. Gurfinkel, V. et al. Postural muscle tone in the body axis of healthy humans. J. Neurophysiol. 96, 2678–2687 (2006).

75. Petzold, B. C. et al. Caenorhabditis elegans body mechanics are regulated by body wall muscle tone. Biophys. J. 100, 1977–1985 (2011).

76. Cacciatore, T. W., Anderson, D. I. & Cohen, R. G. Central mechanisms of muscle tone regulation: implications for pain and performance. Front. Neurosci. 18, 1511783 (2024).

77. Kohsaka, H., Takasu, E., Morimoto, T. & Nose, A. A group of segmental premotor interneurons regulates the speed of axial locomotion in Drosophila larvae. Curr. Biol. 24, 2632–2642 (2014).

78. Mark, B. et al. A developmental framework linking neurogenesis and circuit formation in the Drosophila CNS. Elife 10, (2021).

79. Heckscher, E. S., Lockery, S. R. & Doe, C. Q. Characterization of Drosophila larval crawling at the level of organism, segment, and somatic body wall musculature. J. Neurosci. 32, 12460–12471 (2012).

80. Han, C., Jan, L. Y. & Jan, Y.-N. Enhancer-driven membrane markers for analysis of nonautonomous mechanisms reveal neuron-glia interactions in Drosophila. Proc. Natl. Acad. Sci. U. S. A. 108, 9673–9678 (2011).

81. Pfeiffer, B. D., Truman, J. W. & Rubin, G. M. Using translational enhancers to increase transgene expression in Drosophila. Proc. Natl. Acad. Sci. U. S. A. 109, 6626–6631 (2012).

82. Mahr, A. & Aberle, H. The expression pattern of the Drosophila vesicular glutamate transporter: a marker protein for motoneurons and glutamatergic centers in the brain. Gene Expr. Patterns 6, 299–309 (2006).

83. Fei, H. et al. Mutation of the Drosophila vesicular GABA transporter disrupts visual figure detection. J. Exp. Biol. 213, 1717–1730 (2010).

84. Li, C. H. & Lee, C. K. Minimum cross entropy thresholding. Pattern Recognit. 26, 617–625 (1993).

